# Enhancing bacteriophage therapeutics through *in situ* production and release of heterologous antimicrobial effectors

**DOI:** 10.1101/2022.03.09.483629

**Authors:** Jiemin Du, Susanne Meile, Jasmin Baggenstos, Tobias Jäggi, Pietro Piffaretti, Laura Hunold, Cassandra I. Matter, Lorenz Leitner, Thomas M. Kessler, Martin J. Loessner, Samuel Kilcher, Matthew Dunne

## Abstract

Bacteriophages operate via pathogen-specific mechanisms of action distinct from conventional, broad-spectrum antibiotics and are emerging as promising alternatives. However, phage-mediated killing is often limited by bacterial resistance development (1,2). Here, we engineer phages for target-specific effector gene delivery and host-dependent production of colicin-like bacteriocins and cell wall hydrolases. Using urinary tract infection (UTI) as a model, we show how heterologous effector phage therapeutics (HEPTs) suppress resistance and improve uropathogen killing by dual phage- and effector-mediated targeting. Moreover, we designed HEPTs to control polymicrobial uropathogen communities through production of effectors with cross-genus activity. Using a phage-based companion diagnostic (3), we identified potential HEPT responder patients and treated their urine *ex vivo*. Compared to wildtype phage, a colicin E7-producing HEPT demonstrated superior control of patient *E. coli* bacteriuria. Arming phages with heterologous effectors paves the way for successful UTI treatment and represents a versatile tool to enhance and adapt phage-based precision antimicrobials.

## Main

Currently, conventional, small-molecule antibiotics with broad target specificity are the most effective treatments against bacterial infections. However, the global emergence and spread of antimicrobial resistance (AMR) (4), as well as adverse effects caused by antibiotic-induced microbiome dysbiosis, highlight the need for novel and more pathogen-specific antimicrobial interventions (5,6). Bacteriophages (phages), bacteriocins, synthetic antimicrobial peptides, and target-specific cell wall hydrolases (e.g., phage-derived endolysins) are currently being developed as precision antimicrobials (7). Among these, phages are highly promising because of their ubiquity, pathogen specificity, and ability to self-replicate (8,9). Although the killing of host bacteria by phages is largely independent of the host drug-resistance profile, treatment with phages often fails to inactivate all bacterial cells within a target population. This can be due to phage tolerance (10) or resistance mechanisms that bacteria employ to counteract viral predation, including the production of extracellular matrices, mutation or reduced expression of phage receptors, adaptive CRISPR-Cas immunity, restriction/modification systems, abortive infection systems, and a growing number of other resistance mechanisms described in the literature (11,12).

Recent advances in CRISPR-Cas technology and synthetic biology have enabled the rapid modification of phage genomes beyond model phages (such as T4, T7, or lambda) to include therapeutic phage candidates that are typically less well-studied (13). As a result, engineering has been applied to (i) adapt phage tropism through directed receptor binding protein modification (14–17), (ii) construct sequence-specific antimicrobials through phage-mediated, pathogen-specific delivery of programmed CRISPR-Cas modules (18,19), (iii) deliver toxic proteins as genetic payloads (20), (iv) develop rapid phage-based (companion) diagnostics through the delivery of reporter genes (reporter phages) (13,21), and (v) optimize therapeutic phages for experimental therapy (22).

In this study, we demonstrate how diverse phages can be engineered to encode bacteriocins and cell wall hydrolases as antimicrobial effector genes, a concept we coin <u>h</u>eterologous <u>e</u>ffector <u>p</u>hage <u>t</u>herapeutics (HEPTs). Here, effector genes are expressed during infection and their products released upon host cell lysis to function as secondary pathogen-specific antimicrobials, thereby complementing and enhancing phage-mediated killing. As a model system, we focused on developing HEPTs as precision antimicrobials against UTI pathogens (concept: **Fig. 1a**). UTIs are among the most common community-acquired and healthcare-associated microbial infections in all age groups and a major public health concern, resulting in annual healthcare costs exceeding 1.6 billion dollars in the US alone (23). While the most prevalent causative agent of UTIs is *Escherichia coli*, the microbial etiology is complex and can involve a wide range of Gram-negative or Gram-positive bacteria and certain fungi (24). An analysis of 340 isolates acquired from 231 incidents of UTI during 2020 in Zurich, Switzerland (the Zurich Uropathogen Collection; **Fig. S1**) identified 26 different bacterial species, with *E. coli* (34%), *Enterococcus faecalis* (17%), and *Klebsiella pneumoniae* (14%) as the most prevalent uropathogens, which is consistent with previous etiological studies on UTIs (24).

**Figure 1.**
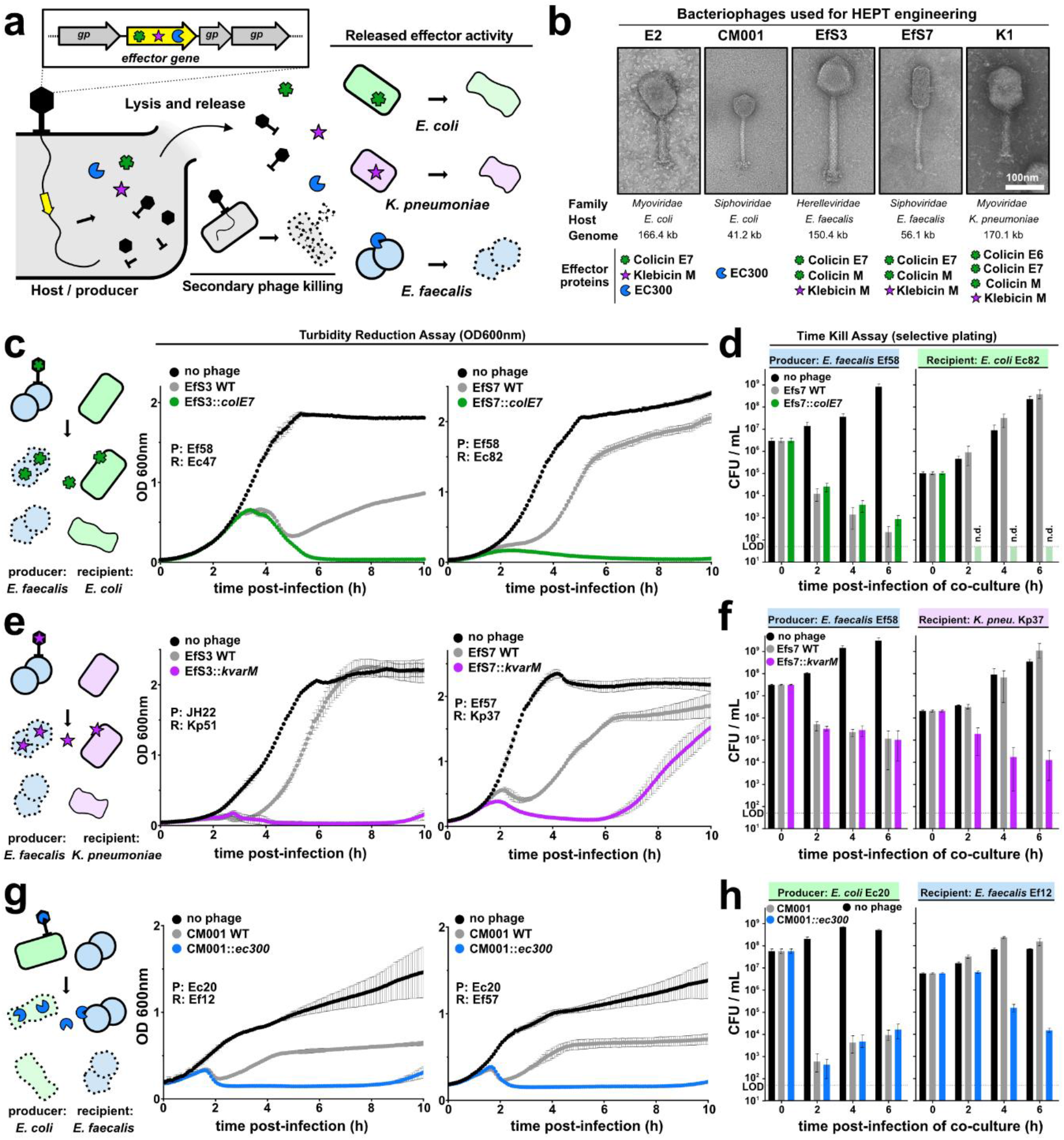
Cross-targeting HEPTs control polymicrobial uropathogen communities. **(a)** Heterologous effector phage therapeutics (HEPTs) enable pathogen-specific delivery and production of antimicrobial effector genes (yellow). Upon phage-induced host cell lysis, effectors are released alongside progeny virions to exert a secondary antimicrobial activity against defined bacterial targets. HEPTs were designed against the uropathogens *E. coli* (light green), *K. pneumoniae* (light purple), and *E. faecalis* (light blue). **(b)** Five phages were employed as HEPT scaffolds to integrate colicins (M, E6, and E7; square star), klebicin M (star), or the *E. faecalis*-specific cell wall hydrolase EC300 (Pac-Man) (28) within the phage structural gene cassette. The colour of the effector symbol matches the target organism **(c-h)** The antimicrobial effect of cross-targeting HEPTs was compared to their WT counterparts upon infecting co-cultures of a phage-susceptible host (P, producer) and an effector-susceptible target (R, recipient). Bacterial killing was quantified using 10 h turbidity reduction assays (**c**, **e**, **g**). 6 h time kill assays were combined with plating on differential and selective agar to enumerate different bacterial species **(d, f, h)**. Strains used from the Zurich Uropathogen Collection: *E. faecalis* Ef12/57/58; *E. coli* Ec20/47/82; *K. pneumoniae* Kp37/51; gp = gene product; LOD = limit of detection; OD_600nm_ = optical density at 600 nm. Data are shown as mean ± SD (n=3).

Guided by these observations, we engineered HEPTs using five distinct and strictly lytic phages that target the predominant uropathogens *E. coli* (phages E2 and CM001), *E. faecalis* (phages EfS3 and EfS7), and *K. pneumoniae* (phage K1). These phages represent various phylogenetic families with distinct virion morphologies and genome sizes (3,25) (**Fig. 1b, Table S1**). To target Gram-negative uropathogens, we selected four colicin-like bacteriocins (CLBs) as effectors, which are active against either *E. coli* (colicins E6, M, and E7; *green*) or *K. pneumoniae* (klebicin M; *purple*). CLBs are protein toxins with narrow-spectrum activity, characterized by a distinct, three-domain architecture that mediates receptor binding (central domain), Ton- or Tol-dependent translocation (N-terminal domain), and periplasmic or intracellular toxicity (C-terminal domain) (26). The cytotoxicity of CLBs that were selected for HEPT engineering is mediated by periplasmic peptidoglycan biosynthesis inhibition (colicin M, klebicin M), intracellular 16s rRNase activity (colicin E6), or unspecific cytosolic nuclease activity (colicin E7) (26,27). To target the Gram-positive uropathogen *E. faecalis*, we employed a phage-derived, chimeric cell wall-hydrolase (EC300; *blue*) that recognizes and degrades *E. faecalis* cell walls with high specificity (28). EC300 was engineered by fusing an M23 endopeptidase domain from a virion-associated lysin with a cell wall binding domain of an *E. faecalis* phage endolysin (28).

All effector genes were codon-optimized to match scaffold target species specificity (29) and integrated within the phage structural gene cassette alongside a strong ribosomal binding site to guide late promoter-driven expression (see **Fig. S2a**). Engineered HEPT candidates were constructed either using CRISPR-Cas9-assisted engineering (3,13,25) or by rebooting synthetic genomes in suitable surrogate hosts (see **online methods**) (30). To avoid toxicity, CLB-encoding HEPTs were engineered and amplified under constitutive expression of their respective immunity proteins. The production of active effector protein upon engineered phage infection was demonstrated by spot assays using crude wildtype (WT) phage or HEPT lysates on a selection of clinical target strains (**Fig. S2b, c, d**). To ensure that phage activity does not interfere with these assays, we tested lysates from engineered phages producing effectors with cross-genus activity. All effectors were produced and active against a broad range of urine-derived isolates of the respective target species with variable levels of activity depending on the phage scaffold (**Tables S2&S3**). The klebicin M and EC300 encoding HEPTs targeted up to 54% and 92% of tested isolates, respectively. Among the three colicins, colicin E7 presented the broadest range of activity with HEPT EfS3::*colE7* active against 70% of the 56 *E. coli* isolates tested.

Polymicrobial infections are commonly observed within the urinary tract, particularly during catheter-associated UTIs (31), which may complicate therapeutic phage selection, combination, and treatment. Interestingly, analysis of the Zurich Uropathogen Collection revealed 34% of UTI cases (78/231) as polymicrobial with *E. faecalis* identified as a common co-infector associated with polymicrobial UTIs involving *E. coli* (46%) and *K. pneumoniae* (39%) (**Fig. S1c**). We therefore assessed the ability of HEPTs to deliver effectors with cross-genus activity to target polymicrobial communities composed of different combinations of clinical isolates through enzymatic collateral damage (**cross-targeting HEPTs**, **Fig. 1c-h**). Using turbidity reduction and time kill assays combined with differential plating, we demonstrate the ability of HEPTs EfS3::*colE7* and EfS7::*colE7* to infect and kill *E. faecalis* cells (P, producer) while simultaneously eradicating co-cultured *E. coli* (R, recipient) within two hours of treatment through *in situ* release of colicin E7 effectors (**Fig. 1c-d**). Similarly, we demonstrate that the klebicin M-producing HEPTs EfS3::*kvarM* and Efs7::*kvarM* can successfully control co-cultures of *E. faecalis* (P) and *K. pneumoniae* (R) (**Fig. 1e-f**), suggesting that CLB effectors provide a viable strategy for cross-genus targeting.

Since the protective peptidoglycan layer of Gram-positive pathogens such as *E. faecalis* is externally accessible, cell wall hydrolases are also promising enzyme antibiotics (enzybiotics) for cross-genus HEPT engineering. As shown in **Fig. 1g-h**, we demonstrate the ability of an EC300-producing HEPT based on *E. coli* phage CM001 (CM001::*ec300*) to target *E. faecalis* in co-culture, resulting in approximately 4-log reduction in *E. faecalis* after six hours of treatment for a co-culture of *E. coli* Ec20 (P) and *E. faecalis* Ef12 (R) (**Fig 1h**). In clinical practice, polymicrobial UTIs would require application of multiple phages for treatment. Using a combination of two cross-genus HEPTs (E2::*kvarM* and K1::*colE7*), we were able to demonstrate a strongly enhanced killing of an *E. coli* and *K. pneumoniae* co-culture, providing a promising strategy to tackle polymicrobial UTIs (**Fig. S3a**). During polymicrobial UTIs, each bacterial species could be leveraged as an effector-producing host, even if it might not be the causative agent.

Regardless of the importance of polymicrobial infections, most UTIs are caused by a single uropathogen, with *E. coli* and *K. pneumoniae* as predominant agents (24). Infection of monocultures with WT phages typically leads to substantial initial host killing, as can be observed in turbidity reduction assays. However, within hours of infection, stable or transient phage resistance frequently occurs, leading to regrowth of phage resistant populations. To demonstrate this well-known limitation for phage treatment, we infected urine-derived *E. coli* (Ec20 and Ec41) or *K. pneumoniae* (Kp18, Kp28, and Kp37) isolates with WT phages E2 or K1, respectively. As expected, regrowth was observed within <18 h and a second round of infection demonstrated that these cells no longer responded to phage challenge, with both transient and stable phage resistance identified for individual clones isolated after the second round of infection (**Fig. S4**).

To circumvent this limitation, HEPTs were engineered to target resistant subpopulations through phage-mediated delivery of CLBs that provide an orthogonal killing mechanism against the same target species (**self-targeting HEPTs**, **Fig. 2a**). To this end, we constructed HEPTs E2::*colE7* and K1::*kvarM* (**Fig. 2b**) to treat *E. coli* or *Klebsiella* monocultures and compared performance to their WT phage counterparts. Depending on the target isolate, E2 WT treatment led to rapid (*E. coli* Ec20) or delayed (*E. coli* Ec41) regrowth due to the expansion of resistant subpopulations. Strikingly, treatment with E2::*colE7* led to a sustained (18 h) reduction in optical density and dramatic reduction of *E. coli* cell counts (>6-log reduction as compared to E2 WT) (**Fig. 2c**). Similar results were obtained for the self-targeting HEPT K1::*kvarM*, which strongly reduced *Klebsiella* cell counts at 10 h post-infection (p.i.) and delayed the regrowth of *Klebsiella* cells when measured at 18 h p.i. (**Fig. 2d**). Akin to the use of dual cross-targeting HEPTs (**Fig. S3a**), the combination of E2::*colE7* and K1::*kvarM* led to enhanced control of *E. coli* and *Klebsiella* co-cultures compared to combinations containing WT phages (**Fig. S3b**). Overall, through *in situ* production of self-targeting effectors, HEPTs provide an effective two-pronged attack to target bacteria and prevent or delay their regrowth.

**Figure 2.**
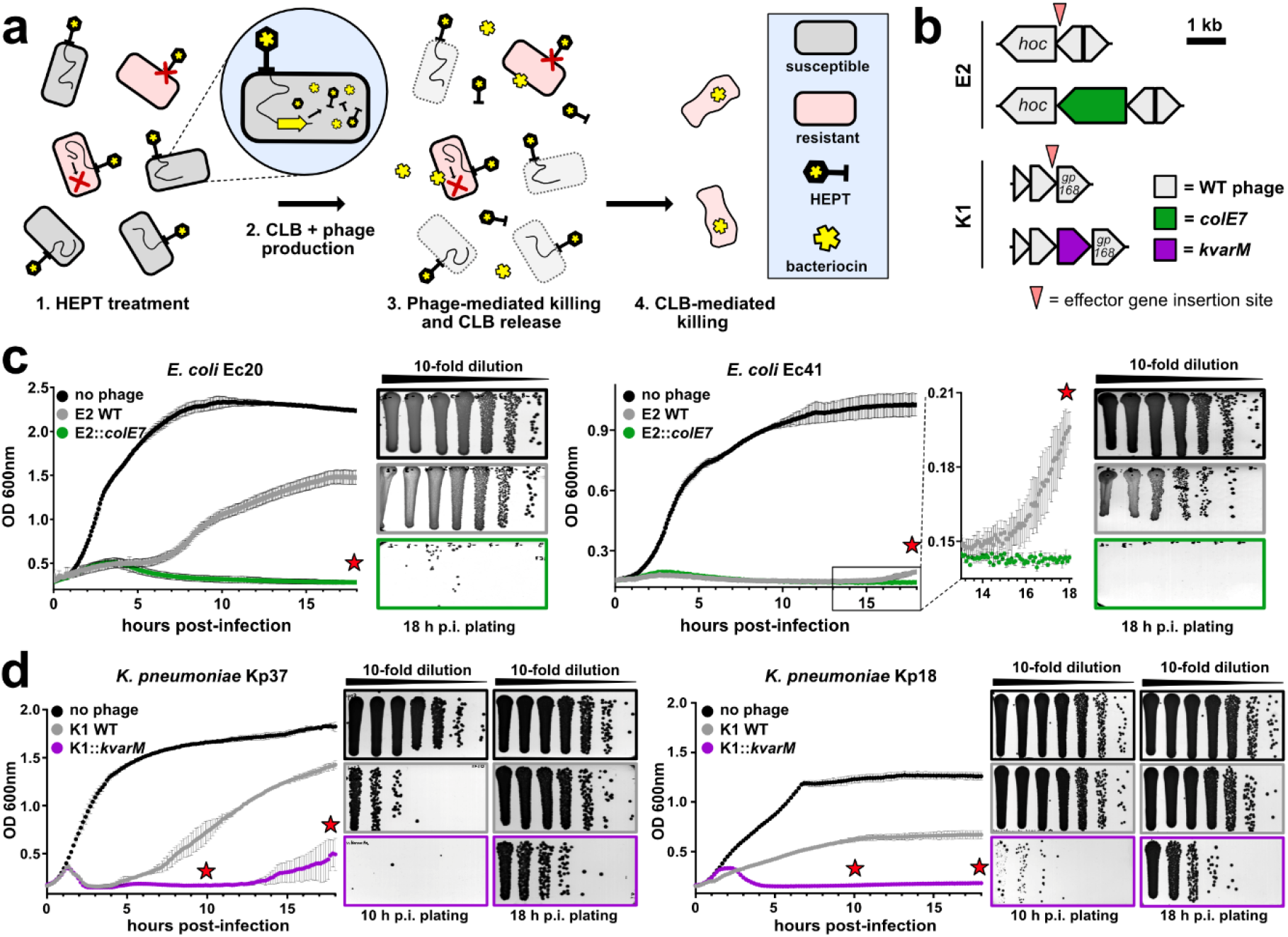
Self-targeting HEPTs enhance killing of uropathogenic *E. coli* and *K. pneumoniae* isolates through release of colicin-like bacteriocins. **(a)** The rapid propagation of bacteria (pink) surviving WT phage treatment (e.g., phage-resistant subpopulations) results in a failure to control bacterial growth. Self-targeting HEPTs can prevent or delay the growth of resistant subpopulations *in vitro* by releasing complementary antimicrobial effectors. **(b)** Genes encoding colicin E7 (green) or klebicin M (purple) were integrated within the structural gene cassette of phage scaffolds E2 or K1 to generate HEPTs targeting *E. coli* (E2::*colE7*) or *K. pneumoniae* (K1::*kvarM*). **(c-d)** Turbidity reduction assays combined with timepoint plating (red stars) demonstrated improved antimicrobial activity (i.e., regrowth was avoided or delayed) for E2::*colE7* **(c)** and K1::*kvarM* **(d)** compared to WT phage treatment of uropathogenic *E. coli* and *K. pneumoniae* monocultures, respectively. *hoc* = highly immunogenic outer capsid protein. HEPTs = heterologous effector phage therapeutics, CLB = colicin-like bacteriocin. OD_600nm_ = optical density at 600 nm. Data are shown as mean ± SD (n=3).

In the future, phage-based precision antimicrobials will most likely be designed and implemented as personalized treatment options. Therefore, a rapid and reliable companion diagnostic would be helpful to guide phage selection and/or predict therapeutic success. To assess the performance of self-targeting HEPTs against *E. coli* in patient urine, we combined our recently developed reporter phage-based diagnostic (3) with *ex vivo* urine treatment using HEPT E2::*colE7* (workflow: **Figure 3a**). To this end, 39 patient urine samples were collected and subjected to reporter phage-based pathogen identification and phage susceptibility screening using a nanoluciferase-encoding phage E2 (E2::*nluc*) (3). Reporter phage-induced bioluminescence was quantified within 4.5 h as an indicator of successful phage genome delivery and expression. Concurrently, the presence of *E. coli* in individual urine specimens was screened and confirmed using differential plating. Eight urine samples were positive for *E. coli*, seven of which were detected by E2::*nluc* (**Fig. 3b**, urinalysis).

**Figure 3.**
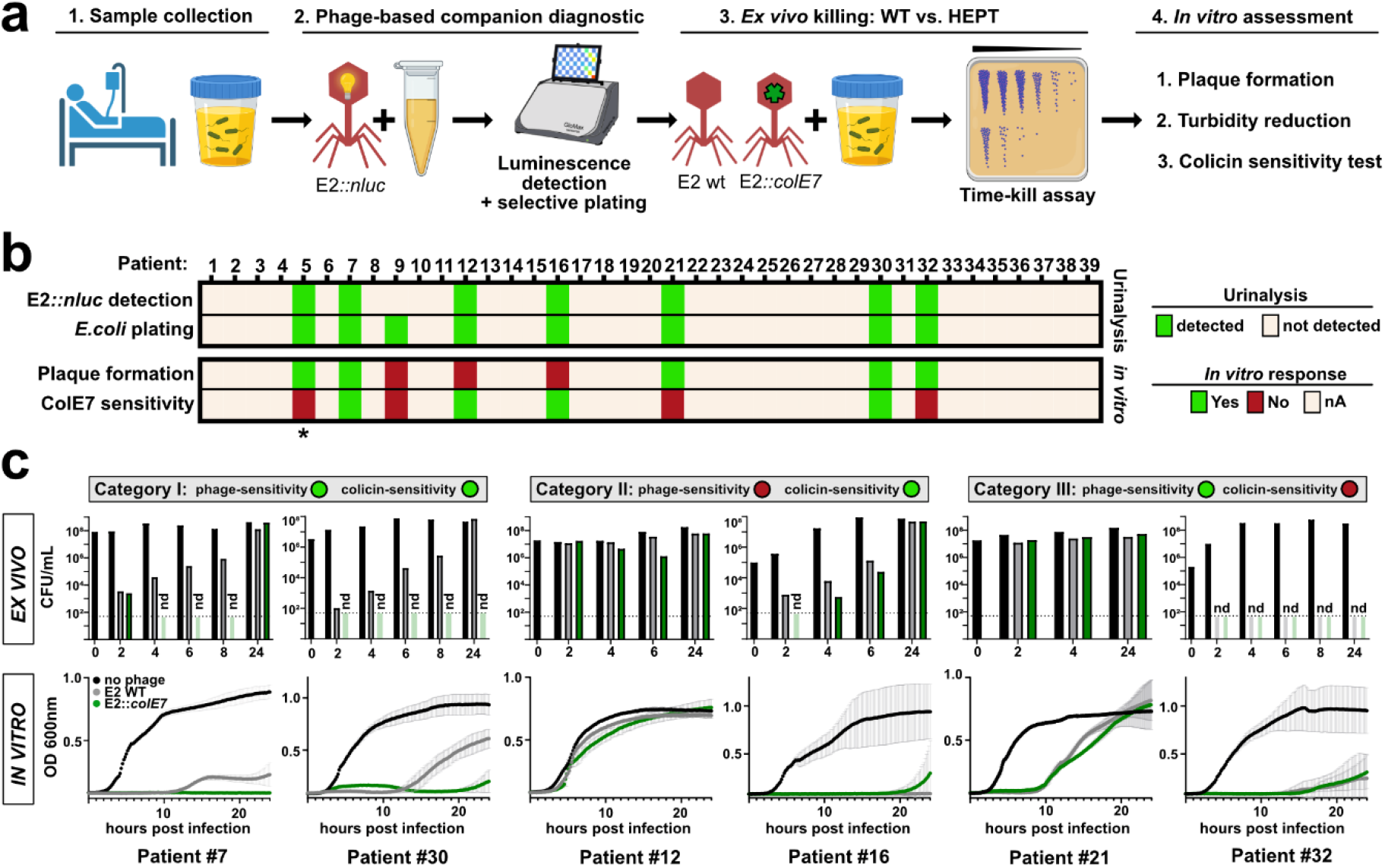
HEPTs provide enhanced killing of colicin-sensitive *E. coli* in patient urine. **(a)** Workflow for combining a phage-based companion diagnostic to identify potential HEPT responder patients presenting with E2-sensitive *E. coli* bacteriuria (**steps 1-2**) with subsequently *ex vivo* treatment (**step 3**) and *in vitro* assessment (**step 4**) of positive urine specimens using E2 WT or E2::*colE7*. **(b)** Patient urine samples (n=39) were subjected to a bioluminescence-based (E2::*nluc*) reporter phage assay (3) to identify E2-sensitive *E. coli* in the urine within 4.5 h (3). Urine was plated on differential agar to isolate patient strains and enumerate overall bacterial load. *E. coli* isolates were further tested *in vitro* for E2 sensitivity (plaque formation) and colicin E7 sensitivity (detailed results provided in **Fig. S5**) and categorized: Category I = phage and colicin sensitive; Category II = phage-resistant, colicin-sensitive; Category III = phage-sensitive, colicin-resistant. **(c)** Time kill assays were used to assess *ex vivo* treatment using 10^9^ PFU/mL E2 or E2::*colE7* added to fresh urine for 24 h at 37 °C. A similar HEPT treatment was performed *in vitro* on patient isolates grown in SHU with bacterial killing measured using turbidity reduction assays. OD_600nm_ = optical density at 600 nm. Turbidity data is mean ± SD (n=3). *, specimen #5 was excluded due to a *K. pneumoniae* polymicrobial infection. Elements of panel **(a)** were created with BioRender.com.

To assess the advantage of heterologous effector delivery, fresh urine samples were subjected to HEPT (E2::*colE7*) or WT phage treatment, with *E. coli* killing assessed over 24 h using time kill assays (**Fig. 3c**, *ex vivo*). In addition, patient-derived *E. coli* strains were isolated and tested for phage and colicin E7 susceptibility, with phage susceptibility defined as the ability to form plaques (**Fig. 3b**, **Fig. S5**). Specimen #5 was excluded due to the presence of >10^7^ CFU/mL *K. pneumoniae* (polymicrobial infection). The remaining six patient isolates could be classified into three categories based on their susceptibility profiles: (I) phage and colicin sensitive, (II) phage resistant but colicin sensitive, and (III) phage sensitive but colicin resistant. Compared to E2 WT, improved *E. coli* killing by E2::*colE7* was observed for both *ex vivo* treatments of urine in category I (specimens #7 and #30), which was subsequently confirmed *in vitro* using turbidity reduction assays (**Fig. 3c**, *in vitro*). In category II (specimens #12 and #16), a slight enhancement of activity was observed during *ex vivo* HEPT treatment; however, no improvement could be validated during *in vitro* turbidity reduction analysis. This was attributed to incomplete infectivity of phage E2 towards specimens #12 and #16, as observed by a lack of plaque formation. As expected, due to a lack of colicin sensitivity for specimens #21 and #32, no difference in activity was observed between E2 WT and E2::*colE7* within category III. Overall, the *ex vivo* study demonstrated enhanced HEPT-mediated killing of *E. coli* in fresh patient urine, provided that the isolate is susceptible to both phage and effector, e.g., colicin E7. For future implementation of HEPTs and other phage-based therapeutics, careful and rapid screening of relevant susceptibility profiles, e.g., using reporter phage companion diagnostics, will be an essential component to guide their therapeutic use.

In conclusion, we present HEPTs as precision antimicrobials that combine the inherent, pathogen-specific killing activity of bacteriophages with *in situ* production and release of secondary antimicrobial effectors. This two-pronged approach enhances the antimicrobial activity of phages, is capable of suppressing outgrowth of phage resistant subpopulations, and can be harnessed to provide cross-genus control of bacterial pathogens using a single HEPT. Through the careful selection of phage scaffolds and heterologous effectors, HEPTs provide a customizable platform for targeted antimicrobial therapy.

## Acknowledgements

We would like to thank all members of the CAUTIphage consortium including Jochen Klumpp, Hendrik Koliwer-Brandl, Jonas Marschall, Shawna McCallin, Vera Neumeier, and Reinhard Zbinden for their scientific input and assistance with organizing clinical samples. We thank the clinical care team at the Department of Neuro-Urology, Balgrist University Hospital (Zurich, Switzerland) for urine specimen collection. We also thank Karin Moelling for scientific discussions, Alexander Harms for providing *E. coli* UTI89 and CFT073, and Leo Meile for providing *E. faecalis* JH2-2. Finally, we acknowledge the ScopeM facility at ETH Zurich for assistance with transmission electron microscopy of phage particles.

## CRediT authorship contribution statement

Conceptualization, J.D., S.M., M.J.L., S.K., M.D.; Methodology, J.D., S.M., S.K., M.D.; Project administration, S.K., M.D.; Supervision, S.K., M.D.; Investigation, J.D., S.M., J.B., T.J., P.P., L.M., C.I.M., S.K., M.D.; Data curation, J.D., S.M., S.K., M.D.; Visualization, S.K.; Writing – original draft, S.K., M.D.; Writing – review and editing, J.D., S.M., L.L., T.M.K., M.J.L., S.K., M.D.; Funding acquisition, L.L., T.M.K., M.J.L., S.K., M.D.; Resources, T.M.K., M.J.L., S.K.

## Ethical approval

All patients gave a general written informed consent, in line with the local ethics committee (Kantonale Ethikkommission Zurich, Switzerland), agreeing for further use of health-related personal data and biological material for research purposes. The study was performed in accordance with the World Medical Association Declaration of Helsinki (32) and conformed with the International Conference on Harmonisation (ICH) Good Clinical Practice (GCP) Guidelines (E6) and the International Organization for Standardization (ISO, 14,155).

