## Supplemental Methods, Figures, and Tables for "Enhancing bacteriophage therapeutics through *in situ* production and release of heterologous antimicrobial effectors"

**<u>Supplemental and Online Material</u>**

**- Online Materials and Methods**

**- Supplementary Figures S1-S5**

**Figure S1**. Analysis of UTI incidents within the Zurich Uropathogen Collection.

**Figure S2**. HEPT construction and assessment of payload activity.

**Figure S3**. Simultaneous application of two cross- or two self-targeting HEPTs to co-cultures containing uropathogenic *E. coli* and *K. pneumoniae*.

**Figure S4**. Phage resistance development upon treatment of UTI isolates with wildtype E2 and K1.

**Figure S5**. Reporter phage-based urinalysis of patient urine samples with phage and colicin E7 susceptibility screening of isolated, patient-derived *E. coli* strains.

- **Supplementary Tables**

**Table S1**. Phages used in the present study and their propagation hosts.

**Table S2**. Bacteriocin payload assessment.

**Table S3**. Peptidoglycan hydrolase (EC300) payload assessment.

**Table S4**. Plasmids and primers used for phage engineering.

**Table S5**. Synthetic DNA strings used for pSelect and pEdit plasmid generation.

**Table S6**. Primers and templates used for synthetic HEPT phage construction.

### Materials and Methods (online)

#### *Bacterial strains and culture conditions*

*E. coli* BL21 (New England Biolabs), *E. coli* Ec20 (1), *K. pneumoniae* KpGe (2), *E. faecalis* JH2-2, and *E. faecalis* Ef57 (3) were used as phage propagation and engineering hosts. *E. coli* XL1-Blue MRF' (Stratagene) was used as a cloning host for plasmid construction. Clinical strains used in this study are listed in **Tables S2 and S3**, which includes isolates taken from the Zurich Uropathogen Collection; a library of 665 patient isolates identified from urine specimens of patients from the Department of Neuro-Urology, Balgrist University Hospital, Zurich, Switzerland acquired between January and December 2020 and provided after routine testing by the Institute of Medical Microbiology (IMM), University of Zurich. Gram-negative bacteria were grown at 37 °C in Lysogeny Broth (LB) or Synthetic Human Urine (SHU (4)). Gram-positive bacteria were cultivated at 37 °C in BHI-fc broth (37 g/L Brain-Heart-Infusion broth from Biolife Italiana, 4 g/L glycine, 3.2 mM L-cysteine HCl, 50 mM Tris, 5 ng/mL choline chloride) under microaerophilic conditions.

#### *Phage propagation and purification*

Phages were propagated on their respective propagation hosts using the soft-agar-overlay method as described previously (1,3). To avoid toxicity, *colE7*-encoding HEPTs E2::*colE7*, K1::*colE7*, and K1::*colE6* were propagated in the presence of their immunity plasmids, pIm_E7 and pIm_E6, respectively (**Tables S1 and S4)**. In brief, after overnight incubation, phage particles were extracted using 5 ml SM buffer per plate (50 mM Tris, pH 7.4, 100 mM NaCl, 8 mM $MgSO_4$) and filter-sterilized (0.2 µm) to obtain crude phage lysates (used for effector susceptibility testing as described below). Phage lysates were further purified and concentrated by PEG precipitation (7% PEG 8000 and 1M NaCl) followed by cesium chloride isopycnic centrifugation and finally dialyzed against a 1000-fold excess of SM buffer.

***Phage genome sequencing***

*E. coli* phage CM001 was isolated from a mixture of wastewaters collected in Switzerland using *E. coli* Ec20 as a host, purified using three sequential rounds of the soft-agar-overlay method, and further propagated as described above. Genomic DNA was extracted from purified phage particles using the phenol:chloroform:isoamyl alcohol (25:24:1) extraction method. Purified DNA was Illumina sequenced (2 x 150bp) by Eurofins Genomics Europe Sequencing GmbH (Constance, Germany). A single contig was obtained by *de novo* assembly using the CLC Genomics Workbench version 20 (QIAGEN Bioinformatics) with default settings. Coding DNA sequence (CDS) identification and annotation was performed using the RAST server (5), with tRNAscan-SE used to identify possible tRNA genes (none detected) (6). Subsequent manual curation and validation was performed using related *E. coli* phage K1H (NC_027994) as a reference genome. The final annotated genome of phage CM001 is available from the GenBank database (UTI-CM001, OM810255) alongside previously sequenced genomes of phage E2 (OL870316), K1 (OL870318), EfS3 (OL870611), and EfS7 (OL870612).

***Transmission electron microscopy***

Phage particles were negatively stained for 20 seconds with 2% uranyl acetate on carbon-coated copper grids (Quantifoil) and observed at 100 kV on a Hitachi HT 7700 equipped with an AMT XR81B Peltier cooled CCD camera (8M pixel) at the ScopeM facility, ETH Zurich.

***CRISPR-Cas9-assisted phage engineering***

All HEPTs based on phages E2, K1, EfS3, and EfS7::*colM*/*kvarM* were constructed using the homologous recombination-based and CRISPR-Cas9-assisted engineering as previously described (1,3). In short, WT phages were propagated in the presence of the respective editing template (pEdit) to enable sequence-specific transgene integration through homologous recombination. WT phages were selectively restricted using a SpyCas9-based counterselection system (pSelect) directed at the flanking homology arms within individual phage genomes. Silent mutations within the protospacer-adjacent motifs (PAMs) on the homology arms of pEdit enable CRISPR-escape and enrichment of engineered phage. When PAM mutation is impossible, multiple silent mutations were introduced within the SpyCas9-targeted seed sequence (12 nucleotides immediately upstream of the PAM) to abrogate CRISPR targeting. All pEdit and pSelect vectors used for phage construction are compiled in **Table S4**; synthetic DNA strings used for effector amplification are shown in **Table S5**.

***Phage genome assembly and rebooting***

EfS7::*colE7* and CM001::*ec300* genomes were assembled *in vitro* from overlapping (~40bp) PCR fragments using the Gibson isothermal method (NEBuilder HiFi DNA assembly master mix, NEB). 20 ng of PCR products per 1 kb of genomic fragment length were used for assembly. Synthetic genomes of *E. faecalis* HEPTs were rebooted through transfection into *L. monocytogenes* Rev2L L-form bacteria as previously described (7). To reboot CM001::*ec300,* 3 µl of assembly mixture was electroporated into 42 µL of electrocompetent *E. coli* XL1-Blue cells at 1.8 kV, 25 µF, 200 Ω using a BTX ECM630 electroporator (BTX Molecular Delivery Systems, MA, USA). 1 mL of SOC medium was supplemented immediately after electroporation and cells were recovered at 37 °C for 4 h with shaking (180 rpm). Subsequently, 10 µl of chloroform was added to assist host lysis and phage release. Following centrifugation at 12,000 x g for 1 min, dilutions of the supernatant were mixed with 200 µL of an overnight culture of *E. coli* Ec20 and 5 mL of molten LC soft agar, layered onto pre-warmed LB plates, and incubated for 16 h at 37 °C. Synthetic DNA strings used for amplification of effector genes are compiled in **Table S5**. Primers and templates used for phage genome fragmentation and effector gene integration are summarized in **Table S6**.

***Effector susceptibility assessment***

400 μL of a log-phase culture of the target bacterial strain was mixed with 10 mL of molten LC soft agar, poured onto a square plate (12 x 12 cm) containing the appropriate growth agar, and dried for 15 min. 10 μL of each sterile-filtered crude phage lysate was spotted on the bacterial lawn, dried, and incubated overnight to visualize the zones of growth inhibition.

#### ***Turbidity reduction assays***

Log-phase cultures were diluted in BHI-fc (*E. faecalis*) or SHU (*E. coli* and *K. pneumoniae*) to an $OD_{600nm}$ of 0.05 - 0.1, distributed into clear, flat-bottom 96-well plates (Bioswisstech) and infected with phages to obtain a final concentration of $5x10^7$ plaque-forming units (PFU)/mL. The plates were sealed with a microplate sealing film (AxygenTM) and $OD_{600nm}$ was quantified every 5 min at 30 °C using a spectrophotometer (SPECTROstar Omega or SPECTROstar Nano, BMG Labtech). Uninfected bacterial dilutions were used as growth controls, and growth medium without bacteria was used as a background/sterility control. All cross-genus HEPT experiments used a ratio of 10:1 producer to recipient cells. Experiments were performed as technical triplicates and reported as mean ± standard deviation (SD). When indicated, triplicate reactions were combined, serially diluted, and plated on agar plates at 10 or 18 h post-infection.

#### ***Time kill assays***

For cross-genus HEPT TKAs, 1 mL of co-culture was infected with $10^9$ or $5x10^7$ PFU/mL of HEPTs derived from EfS7 and CM001 scaffolds, respectively. The ratio of producer cell to recipient cell was always 10:1 with starting concentrations (CFU/mL) provided in **Fig. 1**. Differential and selective plating was used to enumerate the different bacterial species. *E. coli* and *K. pneumoniae* were counted on chromogenic coliform agar (CCA, Biolife italiana) and *E. faecalis* was counted on KFS agar (Biolife italiana) after incubation at 37 and 42 °C for 16 or 48 h, respectively. Experiments were performed as biological triplicates and reported as mean ± SD.

***Repeated phage exposure and resistance development***

*E. coli* strains Ec20 and Ec41 and *K. pneumoniae* strains Kp18, Kp28, and Kp37 were diluted to an $OD_{600nm}$ of 0.1 and infected with $10^8$ PFU/mL of phages E2 or K1 (round I) in SHU. After 18 h of infection, the phage-exposed samples were combined (n=3) and diluted to an $OD_{600nm}$ of 0.1 in fresh SHU and incubated with additional WT phage or media alone for another 18 h (round II). Individual surviving clones were isolated after 36 h by re-streaking on LB agar and tested *in vitro* for phage susceptibility using spot-on-the-lawn assays.

***Reporter-phage based urinalysis and ex vivo activity assessment***

Reporter phage urinalysis was performed as described previously (1). In brief, 1 mL of patient urine sample was directly mixed with 4 mL of LB. Samples were enriched for 1 h at 37 °C with shaking (180 rpm). 50 µL of reporter phage was added to individual 450 µL aliquots of enriched urine ($10^6$ PFU/mL final concentration) and incubated at 37 °C with shaking. LB media spiked with reporter phages alone served as background controls. Bioluminescence measurements were taken at 3 h post infection. Based on manufacturer's instructions, a buffer-reconstituted NLuc substrate (Nano-Glo Luciferase Assay System, Promega) was mixed at a 1:1 ratio with a sample of the infection mixture (40 µL total) in Nunc™ F96 MicroWell™ plates (Thermo Fisher). Bioluminescence was quantified 5 min after substrate addition using a GloMax® Navigator Luminometer (Promega) with 5 s integration and 2 s delay. Relative light units (RLUs) were background corrected by division of the RLU from phage-only controls. Samples producing $>10^3$ RLU fold change (FC) were considered positive. To confirm reporter phage results and to isolate strains, patient urine was plated on differential agar (UriSelect4, BioRad). *E. coli* isolates were tested for phage-sensitivity by determining the efficiency of plating using the soft-agar-overlay method. For *ex vivo* TKA experiments, 250 µL of patient urine was infected with 250 µL of a $10^9$ PFU/mL stock of E2 or E2::*colE7* in PBS solution, incubated for 24 hours at 37 °C, and plated on LB at the indicated time points. T=0 was plated prior to phage addition as an input control. To replicate TKA conditions, *in vitro* turbidity reduction assays were performed by adding 100 µl of a $10^9$ PFU/mL stock solution of E2 or E2::*colE7* to 100 µl of bacterial culture in SHU at a lower starting $OD_{600nm}$ of 0.005 to reproduce bacterial loads in patient urine.

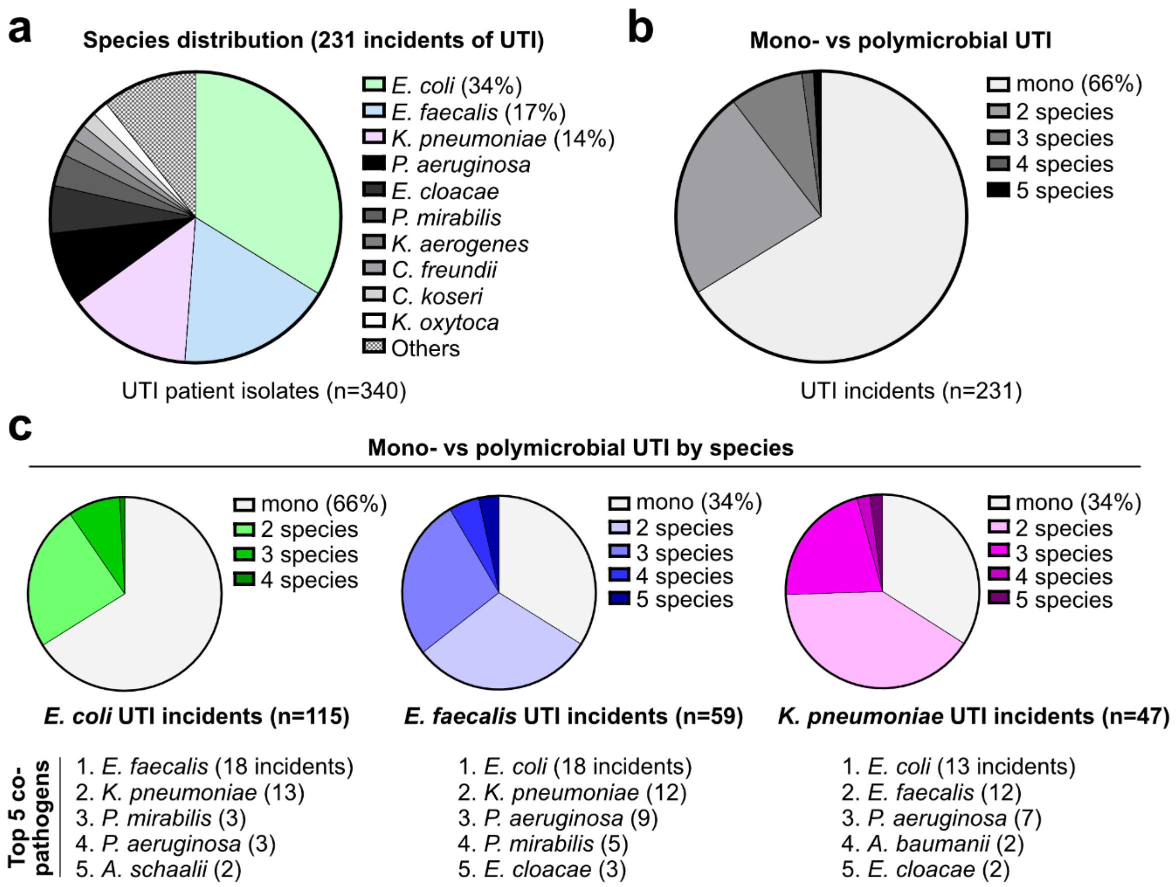


**Figure S1. Analysis of UTI incidents within the Zurich Uropathogen Collection.** The Zurich Uropathogen Collection comprises 665 isolates from 461 incidents of asymptomatic bacteriuria (n=230) or UTI (n=231). (**a**) The species distribution of 340 isolates acquired from UTI patients was determined and (**b**) the occurrence of mono- vs polymicrobial infections quantified for the corresponding UTI incidents. (**c**) UTI incidents involving the top three uropathogens were analyzed separately to determine the top five co-infecting species and the frequency of mono- and polymicrobial UTIs.

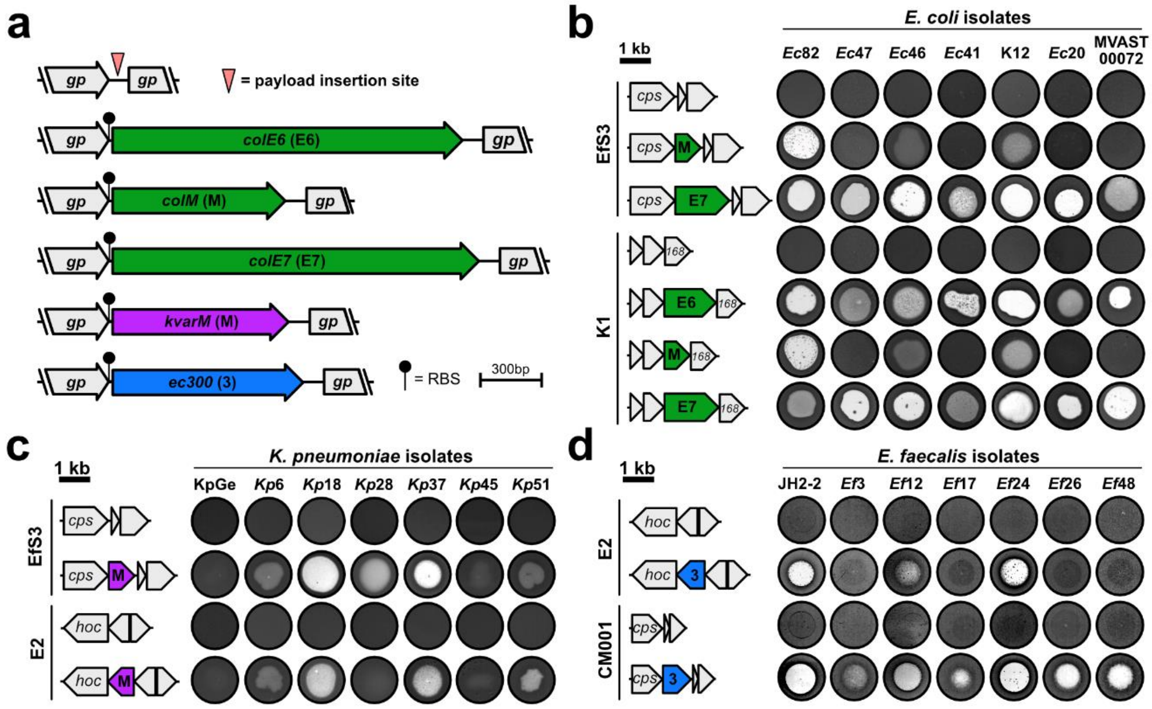


**Figure S2. HEPT construction and assessment of payload activities.** Codon-optimized genes encoding for colicins (M, E6, and E7, green), klebicin M (purple), or the *E. faecalis*-specific cell wall hydrolase EC300 (blue) (8) were integrated within the structural gene cluster of the corresponding phage scaffold alongside a strong ribosomal binding site (RBS) to mediate phage promoter-driven effector expression. Cross-genus antimicrobial activity of crude WT phage or HEPT lysates were tested using the spot-on-the-lawn method against clinical uropathogen isolates (full lists provided in **Tables S2 and S3**). *cps*, major capsid protein; gp, gene product; *hoc*, highly immunogenic outer capsid protein; 168, phage K1 gene product 168; kb, kilobase; HEPTs, heterologous effector phage therapeutics.

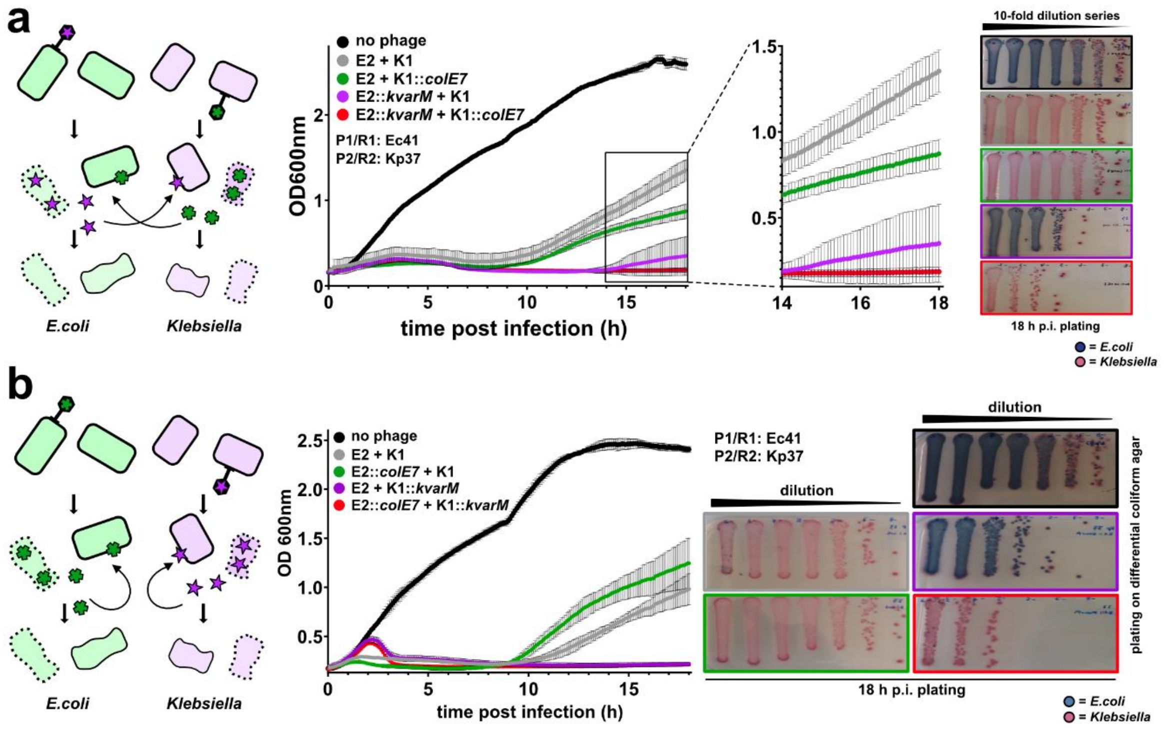


**Figure S3. Simultaneous application of two cross- or two self-targeting HEPTs to co-cultures containing uropathogenic *E. coli* and *K. pneumoniae*.** Growing cultures of *E. coli* (Ec41) and *K. pneumoniae* (Kp37) were adjusted to $OD_{600nm}$ of 0.1, mixed at a ratio of 1:1, and infected with the indicated WT phages and/or HEPTs (5x$10^7$ PFU/mL). Optical density was monitored over 18 h of infection at 30 °C, followed by differential plating on chromogenic coliform agar (matching box and curve colors). Double cross-targeting with phages E2::*kvarM* and K1::*colE7* was assessed in **(a)**, while double self-targeting with phages E2::*colE7* and K1::*kvarM* was assessed in **(b)**. P, producer; R, recipient; HEPTs, heterologous effector phage therapeutics. Data is mean ± SD (n=3).

153

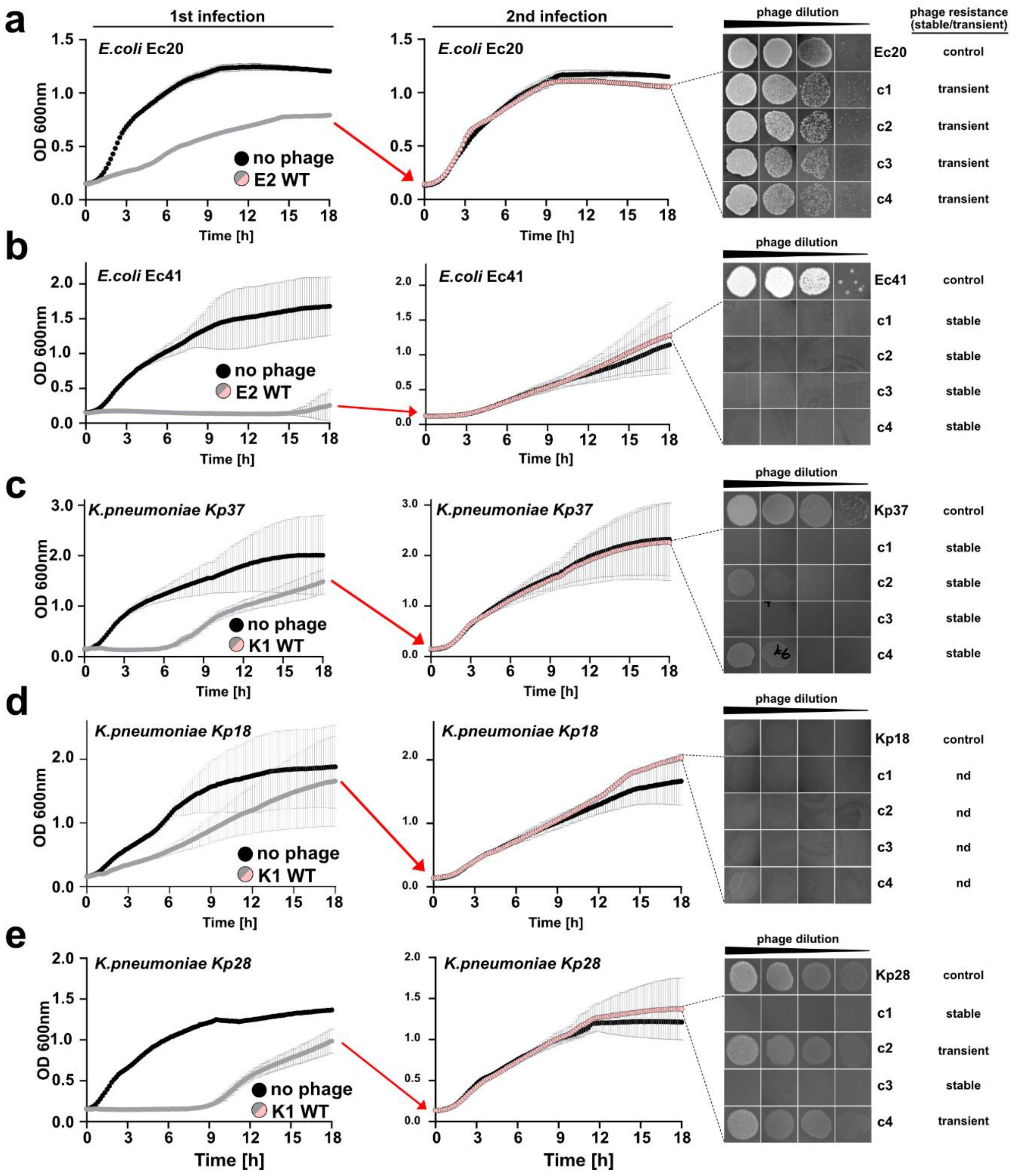


**Figure S4. Phage resistance development upon treatment of UTI isolates with wildtype E2 and K1.** Phage resistance development was assessed by two consecutive rounds of phage infection of bacterial cultures in SHU medium. Turbidity reduction assays were performed for *E. coli* isolates Ec20 **(a)** and Ec41 **(b)** or *K. pneumoniae* isolates Kp37 **(c)** and Kp18 **(d)** infected with $10^8$ PFU/mL of phages E2 or K1, respectively, with optical density monitored for 18 h. Phage-exposed cultures were combined (n=3), re-adjusted to $OD_{600nm}$ of 0.1 in SHU and incubated for another 18 h with additional wildtype (WT) phages (pink) or media alone (black). Growth kinetics were compared to non-infected controls. After the second round of infection, individual clonal survivors were isolated (three rounds of colony purification) and assessed for phage susceptibility using spot-on-the-lawn assays. Turbidity data is mean ± SD (n=3). nd = not determined.

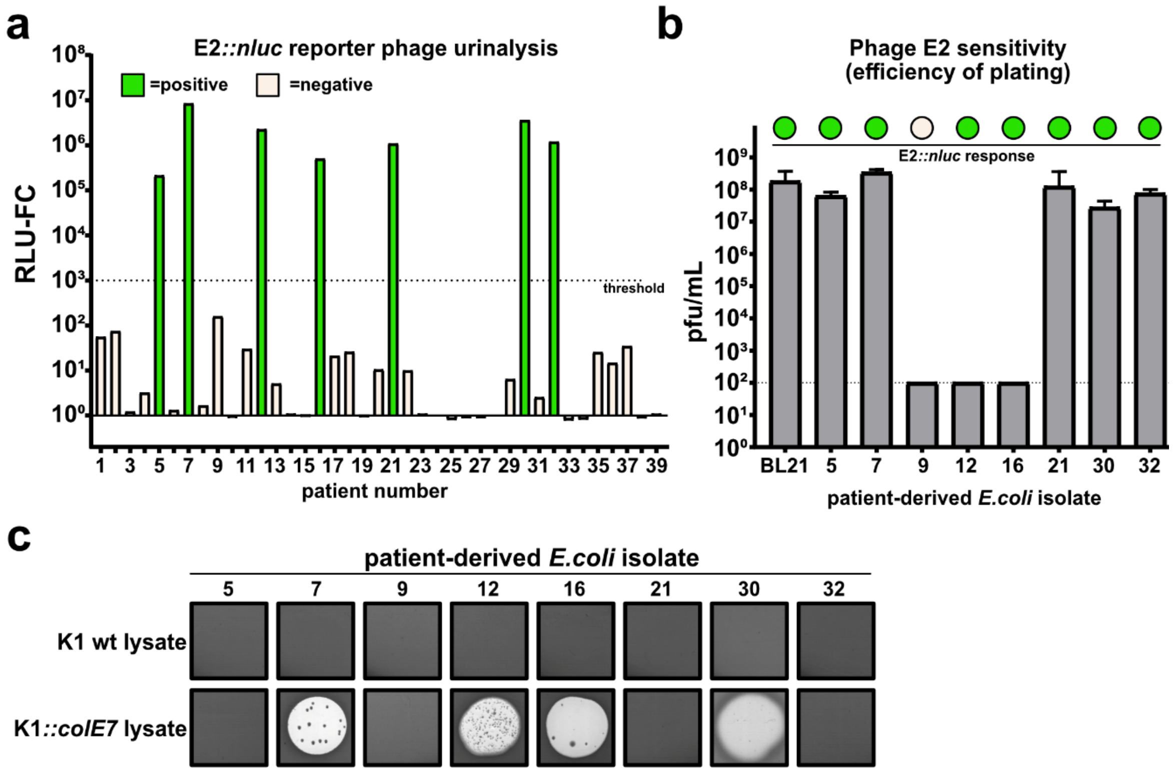


**Figure S5. Reporter phage-based urinalysis of patient urine samples with phage and colicin E7 susceptibility screening of isolated, patient-derived *E. coli* strains. (a)** Reporter phage urinalysis was performed using 39 fresh patient urine samples from the Balgrist University Hospital, Zurich, Switzerland, as described in the **online methods** and in (1). The fold change (compared to media alone) in relative light units (RLU-FC) was determined at 3 hours post-infection with urine samples producing values $>10^3$ considered as positive samples, i.e., containing E2-susceptible *E. coli*. **(b)** All *E. coli* strains were isolated and purified from patient urine after differential plating, and E2-susceptibility quantified using plaque assays (efficiency of plating). **(c)** Colicin E7 effector susceptibility was determined on patient-derived *E. coli* strains by spotting K1 WT or K1::*colE7* phage lysates.

| Phage | Taxonomic classification (GenBank #) | Genome Size [bp] | Payload Target | Payload Nature | Propagation Host | Source |
|---|---|---|---|---|---|---|
| E2 WT | *Caudoviricetes* ; *Caudovirales* ; *Myoviridae; Tevenvirinae; Tequatrovirus* (UTI-E2; OL870316) | 166'367 | - | - | *E. coli* BL21 | [1] |
| E2::*kvarM* | | 167'219 | *Klebsiella spp.* | colM-like murein synthesis inhibitor | *E. coli* BL21 | this study |
| E2::*colE7* | | 168'119 | *E. coli* | unspecific cytosolic nuclease | *E. coli* BL21 (pIm_immE7) | this study |
| E2::*ec300* | | 167'291 | *Enterococcus* | chimeric cell wall-hydrolase | *E. coli* BL21 | this study |
| E2::*nluc* | | 166'904 | *E. coli* | Nanoluciferase | *E. coli* BL21 | [1] |
| K1 WT | *Caudoviricetes; Caudovirales; Myoviridae; Tevenvirinae; Jiaodavirus* (UTI-E2; OL870316) | 170'051 | - | - | *K. pneumoniae* KpGe | [1] |
| K1::*kvarM* | | 170'903 | *Klebsiella spp.* | colM-like murein synthesis inhibitor | *K. pneumoniae* KpGe | this study |
| K1::*colE6* | | 171'728 | *E. coli* | 16s rRNase | *K. pneumoniae* KpGe (pIm_immE6) | this study |
| K1::*colE7* | | 171'803 | *E. coli* | unspecific cytosolic nuclease | *K. pneumoniae* KpGe (pIm_immE7) | this study |
| K1::*colM* | | 170'888 | *E. coli* | murein synthesis inhibitor | *K. pneumoniae* KpGe | this study |
| CM001 WT | *Caudoviricetes; Caudovirales; Siphoviridae; Guernseyvirinae; Kagunavirus* (UTI-CM001; submission OM810255) | 41'222 | - | - | *E. coli* Ec20 | this study |
| CM001::*ec300* | | 42'146 | *Enterococcus* | chimeric cell wall-hydrolase | *E. coli* Ec20 | this study |
| Efs3 WT | *Caudoviricetes; Caudovirales; Herelleviridae; Brockvirinae* (UTI-EfS3; OL870611) | 150'393 | - | - | *E. faecalis* JH2-2 | [3] |
| Efs3::*colE7* | | 152'140 | *E. coli* | unspecific cytosolic nuclease | *E. faecalis* JH2-2 | this study |
| Efs3::*colM* | | 151'225 | *E. coli* | murein synthesis inhibitor | *E. faecalis* JH2-2 | this study |
| EfS3::*kvarM* | | 151'240 | *Klebsiella spp.* | colM-like murein synthesis inhibitor | *E. faecalis* JH2-2 | this study |
| Efs7 WT | *Caudoviricetes; Caudovirales; Siphoviridae; Saphexavirus* (UTI-EfS7; OL870612) | 56'144 | - | - | *E. faecalis* Ef57 | [3] |
| Efs7::*colE7* | | 57'891 | *E. coli* | unspecific cytosolic nuclease | *E. faecalis* Ef57 | this study |
| Efs7::*colM* | | 56'976 | *E. coli* | murein synthesis inhibitor | *E. faecalis* Ef57 | this study |
| Efs7::*kvarM* | | 56'991 | *Klebsiella spp.* | colM-like murein synthesis inhibitor | *E. faecalis* Ef57 | this study |

**Table S1. Phages used in the present study and their propagation hosts.**

| | | | *Enterococcus* phages | | | | *Klebsiella* phage | | |
|---|---|---|---|---|---|---|---|---|---|
| **Species** | **Designation** | **Source** | **EfS3::colM** | **EfS3::colE7** | **EfS7:colM** | **EfS7::colE7** | **K1::*colM*** | **K1::*colE7*** | **K1::*colE6*** |
| *E. coli* | BL21 | 1 | - | + | - | + | - | - | - |
| *E. coli* | K-12 | 2 | + | ++ | + | ++ | + | ++ | ++ |
| *E. coli* | XL1-Blue (K-12 derivative) | 2 | + | ++ | + | ++ | + | ++ | ++ |
| *E. coli* | CFT073 | 3 | - | - | - | - | - | - | + |
| *E. coli* | UTI89 | 3 | + | + | + | + | + | + | - |
| *E. coli* | MVAST-0072 | 4 | - | ++ | - | ++ | - | | ++ |
| *E. coli* | JJ1887 | 4 | - | + | - | + | - | + | - |
| *E. coli* | Ec3 | 6 | + | + | + | + | + | - | - |
| *E. coli* | Ec22 | 6 | - | - | - | - | - | - | - |
| *E. coli* | Ec42 | 6 | - | - | - | - | - | - | - |
| *E. coli* | Ec43 | 6 | - | ++ | - | ++ | - | ++ | ++ |
| *E. coli* | Ec16 | 6 | - | + | - | + | - | - | + |
| *E. coli* | Ec32 | 6 | - | - | - | - | - | - | - |
| *E. coli* | Ec46 | 6 | + | ++ | + | ++ | + | ++ | ++ |
| *E. coli* | Ec57 | 6 | - | - | - | - | - | - | - |
| *E. coli* | Ec14 | 6 | + | - | - | - | - | - | - |
| *E. coli* | Ec33 | 6 | - | + | - | - | - | - | - |
| *E. coli* | Ec47 | 6 | + | ++ | - | ++ | - | ++ | ++ |
| *E. coli* | Ec54 | 6 | - | ++ | - | ++ | - | ++ | ++ |
| *E. coli* | Ec20 | 6 | + | ++ | + | ++ | + | ++ | ++ |
| *E. coli* | Ec34 | 6 | + | - | - | - | + | - | - |
| *E. coli* | Ec41 | 6 | - | ++ | - | ++ | - | ++ | ++ |
| *E. coli* | Ec62 | 6 | - | + | - | + | - | - | + |
| *E. coli* | Ec28 | 6 | + | ++ | + | ++ | + | - | ++ |
| *E. coli* | Ec52 | 6 | - | + | - | + | - | + | + |
| *E. coli* | Ec63 | 6 | - | - | - | - | - | - | - |
| *E. coli* | Ec65 | 6 | + | + | - | - | - | - | - |
| *E. coli* | Ec56 | 6 | + | ++ | - | ++ | - | ++ | ++ |
| *E. coli* | Ec79 | 6 | - | ++ | - | ++ | - | + | ++ |
| *E. coli* | Ec77 | 6 | - | - | - | - | - | - | - |
| *E. coli* | Ec76 | 6 | + | + | + | + | + | + | + |
| *E. coli* | Ec74 | 6 | + | - | + | - | + | - | - |
| *E. coli* | Ec70 | 6 | - | - | - | - | - | - | - |
| *E. coli* | Ec92 | 6 | - | ++ | - | ++ | - | - | ++ |
| *E. coli* | Ec93 | 6 | + | ++ | + | ++ | + | + | ++ |
| *E. coli* | Ec96 | 6 | - | - | - | - | - | - | - |
| *E. coli* | Ec97 | 6 | - | - | - | - | - | - | - |
| *E. coli* | Ec82 | 6 | ++ | ++ | ++ | ++ | ++ | ++ | ++ |
| *E. coli* | Ec101 | 6 | + | - | - | - | - | - | - |
| *E. coli* | Ec131 | 6 | - | - | - | - | - | - | - |
| *E. coli* | Ec132 | 6 | + | + | - | - | ++ | - | - |
| *E. coli* | Ec136 | 6 | - | ++ | - | ++ | - | ++ | ++ |
| *E. coli* | Ec138 | 6 | + | + | - | + | - | - | - |
| *E. coli* | Ec142 | 6 | + | ++ | + | ++ | - | ++ | ++ |
| *E. coli* | Ec137 | 6 | + | ++ | + | ++ | + | + | ++ |
| *E. coli* | Ec139 | 6 | + | + | - | + | - | - | - |
| *E. coli* | Ec141 | 6 | + | + | + | + | + | + | + |
| *E. coli* | Ec140 | 6 | - | + | - | + | - | - | - |
| *E. coli* | Ec143 | 6 | - | - | - | - | - | - | - |
| *E. coli* | Ec145 | 6 | + | ++ | + | ++ | - | ++ | ++ |
| *E. coli* | Ec146 | 6 | + | ++ | - | ++ | - | ++ | ++ |
| *E. coli* | Ec147 | 6 | + | ++ | + | ++ | + | ++ | ++ |
| *E. coli* | Ec148 | 6 | - | - | - | - | - | - | - |
| *E. coli* | Ec149 | 6 | - | ++ | - | ++ | - | + | ++ |
| *E. coli* | Ec151 | 6 | - | ++ | - | ++ | - | + | ++ |
| *E. coli* | Ec158 | 6 | + | + | - | - | - | - | - |
| | | **Activity** | 27/56 | 39/56 | 16/56 | 35/56 | 15/56 | 26/56 | 29/56 |
| | | **Activity (%)** | 48.2 % | 69.6 % | 28.6 % | 62.5 % | 26.8 % | 46.4 % | 51.8 % |

| | | | *Enterococcus* phages | | | | *Klebsiella* phage | | |
|---|---|---|---|---|---|---|---|---|---|
| **Species** | **Designation** | **Source** | **EfS3::colM** | **EfS3::colE7** | **EfS7:colM** | **EfS7::colE7** | **K1::*colM*** | **K1::*colE7*** | **K1::*colE6*** |
| *E. coli* | Ec20 pIm_immE6 | this study | ++ | ++ | + | ++ | + | ++ | - |
| *E. coli* | Ec20 pIm_immE7 | this study | + | - | + | - | + | - | ++ |

| | | | *Enterococcus* phages | | *E. coli* phage |
|---|---|---|---|---|---|
| **Species** | **Designation** | **Source** | **EfS3::kvarM** | **EfS7::kvarM** | **E2::*kvarM*** |
| *K. pneumoniae* | KpGe | 5 | + | + | + |
| *K. pneumoniae* | 0097 | 4 | - | - | - |
| *K. pneumoniae* | Kp3 | 6 | + | + | + |
| *K. pneumoniae* | Kp37 | 6 | ++ | ++ | ++ |
| *K. pneumoniae* | Kp31 | 6 | - | - | - |
| *K. pneumoniae* | Kp39 | 6 | - | - | - |
| *K. pneumoniae* | Kp5 | 6 | - | - | - |
| *K. pneumoniae* | Kp21 | 6 | - | - | - |
| *K. pneumoniae* | Kp43 | 6 | + | + | + |
| *K. pneumoniae* | Kp16 | 6 | - | - | - |
| *K. pneumoniae* | Kp34 | 6 | - | - | - |
| *K. pneumoniae* | Kp48 | 6 | - | - | - |
| *K. pneumoniae* | Kp45 | 6 | + | + | + |
| *K. pneumoniae* | Kp26 | 6 | - | - | - |
| *K. pneumoniae* | Kp8 | 6 | - | - | - |
| *K. pneumoniae* | Kp22 | 6 | - | - | - |
| *K. pneumoniae* | Kp51 | 6 | + | + | + |
| *K. pneumoniae* | Kp36 | 6 | + | + | + |
| *K. pneumoniae* | Kp6 | 6 | + | + | + |
| *K. pneumoniae* | Kp66 | 6 | - | - | - |
| *K. pneumoniae* | Kp69 | 6 | + | - | - |
| *K. pneumoniae* | Kp11 | 6 | ++ | ++ | ++ |
| *K. pneumoniae* | Kp13 | 6 | ++ | + | + |
| *K. pneumoniae* | Kp14 | 6 | + | + | + |
| *K. pneumoniae* | Kp15 | 6 | - | - | - |
| *K. pneumoniae* | Kp59 | 6 | + | + | + |
| *K. pneumoniae* | Kp70 | 6 | - | - | - |
| *K. pneumoniae* | Kp72 | 6 | - | - | - |
| *K. pneumoniae* | Kp71 | 6 | ++ | + | - |
| *K. pneumoniae* | Kp67 | 6 | + | - | - |
| *K. pneumoniae* | Kp82 | 6 | ++ | + | + |
| *K. pneumoniae* | Kp83 | 6 | + | + | - |
| *K. pneumoniae* | Kp80 | 6 | - | - | - |
| *K. pneumoniae* | Kp81 | 6 | - | - | - |
| *K. pneumoniae* | Kp76 | 6 | ++ | + | + |
| *K. pneumoniae* | Kp78 | 6 | + | + | - |
| *K. pneumoniae* | Kp79 | 6 | - | - | - |
| *K. pneumoniae* | Kp90 | 6 | - | - | - |
| *K. pneumoniae* | Kp93 | 6 | + | - | - |
| *K. pneumoniae* | Kp88 | 6 | + | + | - |
| *K. pneumoniae* | Kp89 | 6 | - | - | - |
| *K. pneumoniae* | Kp94 | 6 | - | - | - |
| *K. pneumoniae* | Kp98 | 6 | + | - | - |
| *K. pneumoniae* | Kp96 | 6 | ++ | + | + |
| *K. pneumoniae* | Kp95 | 6 | - | - | - |
| *K. pneumoniae* | Kp97 | 6 | ++ | + | + |
| *K. pneumoniae* | Kp125 | 6 | + | - | - |
| *K. pneumoniae* | Kp100 | 6 | + | - | - |
| *K. pneumoniae* | Kp20 | 6 | ++ | + | + |
| *K. pneumoniae* | Kp41 | 6 | - | - | - |
| *K. pneumoniae* | Kp46 | 6 | + | + | + |
| *K. pneumoniae* | Kp121 | 6 | - | - | - |
| *K. pneumoniae* | Ko75 | 6 | ++ | ++ | + |
| *K. pneumoniae* | Kp18 | 6 | ++ | ++ | ++ |
| *K. pneumoniae* | Kp28 | 6 | ++ | + | + |
| *K. oxytoca* | Ko101 | 6 | + | - | - |
| *K. oxytoca* | Ko38 | 6 | - | - | - |
| *K. oxytoca* | Ko68 | 6 | - | - | - |
| *K. oxytoca* | Ko87 | 6 | - | - | - |
| | | **Activity** | 32/59 | 25/59 | 21/59 |
| | | **Activity (%)** | 54.20% | 42.40% | 35.60% |

**Table S2. Bacteriocin payload assessment.** 10 µL of phage lysate was spotted on bacterial lawns of the following isolates with activity assessed visually after 16 h incubation at 37°C. ++, complete lysis (clear zone); +, moderate lysis (turbid zone); -, no visible activity. Sources: 1, New England Biolabs, MA, USA (Catalog# C2530H); 2, lab stock; 3, gift from Alexander Harms, University of Basel, Switzerland; 4, BEI Resources, VA, USA (www.beiresources.org); 5, Dicty Stock Center, Northwestern University in Chicago, IL, USA; 6, Zurich Uropathogen Collection (2020).

| *E.faecalis* strains | | | *E. coli* phages | |
|---|---|---|---|---|
| **Species** | **Designation** | **Source** | **E2::*ec300*** | **CM001::*ec300*** |
| *E. faecalis* | JH2-2 | 1 | ++ | ++ |
| *E. faecalis* | Efs3 | 2 | + | ++ |
| *E. faecalis* | Efs12 | 2 | + | ++ |
| *E. faecalis* | Efs17 | 2 | + | ++ |
| *E. faecalis* | Efs26 | 2 | + | ++ |
| *E. faecalis* | Efs29 | 2 | ++ | ++ |
| *E. faecalis* | Efs38 | 2 | + | + |
| *E. faecalis* | Efs48 | 2 | + | ++ |
| *E. faecalis* | Efs49 | 2 | + | ++ |
| *E. faecalis* | Efs57 | 2 | + | ++ |
| *E. faecalis* | Efs58 | 2 | + | ++ |
| *E. faecalis* | Efs73 | 2 | ++ | ++ |
| *E. faecalis* | Efs90 | 2 | - | - |

**Table S3. Peptidoglycan hydrolase (EC300) payload assessment.** 10 µL of phage lysate was spotted on bacterial lawns of the following isolates and activity assessed visually after 16 h incubation at 37°C. ++, complete lysis (clear zone); +, moderate lysis (turbid zone); -, no visible activity. Sources: 1, gift from Leo Meile, ETH Zurich, Switzerland; 2, Zurich Uropathogen Collection (2020).

| **CRISPR-Cas based counterselection plasmids** | | | | | | |
|---|---|---|---|---|---|---|
| **Plasmid name** | **Backbone** | **Plasmid function** | **Spacer sequence (5'-3')** | **Cloning primers** | **Primer Sequence (5'-3')** | **Source** |
| pSelect-E2 | pCas9 (Cam$^R$) | Counterselection of wild-type E2 phage DNA | 1. TTAATAGTCTTTTGACCGGC | pCas9_Fw | TCAGCTAGACTTCAGTCTTGAAAAG | This study |
| | | | 2. GTCTGATCTATAACTCCAGT | pCas9_Bw | GACTCCATTCAACATTGCCGA | |
| pSelect-K1 | pCas9 (Cam$^R$) | Counterselection of wild-type K1 phage DNA | 1. CTGCGTCTACACCAACAGTC | E/K_spacers_Fw | GCAGTAATACAGGGGCTTTTC | This study |
| | | | 2. GTGAGATTCTGCCAGAGCTA | E/K_spacers_Bw | CCTCTTTCTCAAGTTATCATCGG | |
| pSelect-UTI-EfS3 | pLEB579 (Em$^R$) | Counterselection of wild-type EfS3 phage DNA | 1. AACGTGCAATCTTCATCAACAAGGTAGCTG | n/a | n/a | [3] |
| | | | 2. AGACTTATAATATTTACCAAGATGGAGCGA | n/a | n/a | |
| pSelect-UTI-EfS7 | pLEB579 (Em$^R$) | Counterselection of wild-type EfS7 phage DNA | 1. AATTCACCTACGAATACGTCAGCAGTTTCA | n/a | n/a | [3] |
| | | | 2. TAAAATCTACAGTATTACAAAATAAAAAAG | n/a | n/a | |

| **Editing plasmids for homologous recombination-based transgene integration** | | | | | |
|---|---|---|---|---|---|
| **Plasmid name** | **Backbone** | **Insert (sequences available, Table S2)** | **Function** | **Cloning primers** | **Primer Sequence (5'-3')** |
| pEdit_E2 | pUC19 (Amp$^R$) | E2_homology arm synthetic gene string | Vector scaffold for phage E2-based editing template construction. Contain homology arms flanking the designated insertion site (upstream of phage E2 *hoc* ) | pUC19_E2_Fw | GCCTGGGGTGCCTAATGA |
| | | | | pUC19_E2_Bw | TAAGCCAGCCCCGACAC |
| | | | | E2_homology arm_ Fw | AGCTCACTCATTAGGCACCCCAGGCCGTAGCGTTAGTTGCTTCG |
| | | | | E2_homology arm_ Bw | GTTGGCGGGTGTCGGGGCTGGCTTAAATATGACGATGAAGAAGTCAATCC |
| pEdit_E2_kvarM | pEdit_E2 (Amp$^R$) | E/K_kvarM synthetic gene string | Editing template for site-directed *kvarM* insertion upstream of phage E2 *hoc* | pEdit_E2_Fw* | GGATAACTTATGGCTTTTAC |
| | | | | pEdit_E2_Bw* | TTATTTTAATGTTACGAAAGAAG |
| | | | | E_kvarM_Fw | CAACTGTAAAAGCCATAAGTTATCCTTATTTTTTACCAGAACCAGA |
| | | | | E_payload_Bw | CTCTTCTTTCGTAACATTAAAATAAAGTACGAGGAGGTAAATATAT |
| pEdit_E2_colE7 | pEdit_E2 (Amp$^R$) | E/K_colE7 synthetic gene string | Editing template for site-directed *colE7* insertion upstream of phage E2 *hoc* | E_colE7_Fw | CAACTGTAAAAGCCATAAGTTATCCTCATTTACCACGGTGGAT |
| | | | | E_payload_Bw | CTCTTCTTTCGTAACATTAAAATAAAGTACGAGGAGGTAAATATAT |
| pEdit_E2_ec300 | pEdit_E2 (Amp$^R$) | E/K_ec300 synthetic gene string | Editing template for site-directed *EC300* insertion upstream of phage E2 *hoc* | E_ec300_Fw | CAACTGTAAAAGCCATAAGTTATCCTTAAGATTTTTTGGTGATACC |
| | | | | E_payload_Bw | CTCTTCTTTCGTAACATTAAAATAAAGTACGAGGAGGTAAATATAT |
| pIm_immE7 | pRSFDuet-1 (Kan$^R$) | *immE7* synthetic gene string | Plasmid modified for consitutive expression of immunity protein ImmE7 to assist generation of *colE7* engineered phages. | pIm_Fw* | AGCTGTTTCCTGTGTGAAATTC |
| | | | | pIm_Bw* | TCGAGTCTGGTAAAGAAACCG |
| | | | | immE7_Fw | GTGGAATTTCACACAGGAAACAGCTATGGAACTGGAAAACTCTATCTC |
| | | | | immE7_Bw | GCAGCGGTTTCTTTACCAGACTCGATTAACCCTGTTTGAAACCCG |
| pIm_immE6 | pRSFDuet-1 (Kan$^R$) | *immE6* synthetic gene string | Plasmid modified for consitutive expression of immunity protein ImmE6 to assist generation of *colE6* engineered phages. | immE6_Fw | GTGGAATTTCACACAGGAAACAGCTATGGGTCTGAAACTGCACATC |
| | | | | immE6_Bw | GCAGCGGTTTCTTTACCAGACTCGATTACCAGTCACCGTCACGG |
| pEdit_K1 | pUC19 (Kan$^R$) | K1_homology arm synthetic gene string | Vector scaffold for phage K1-based editing template construction. Contain homology arms flanking the designated insertion site (upstream of phage K1 *gp168* ) | pUC19_K1_Fw | GGTGGATGGTTTAATTCTCGCCTGGGGTGCCTAATGA |
| | | | | pUC19_K1_Bw | GCTGCTTCTAAATACTTTGACTAAGCCAGCCCCGACAC |
| | | | | K1_homology arm_ Fw | CACTCATTAGGCACCCCAGGCGAGAATTAAACCATCCACCTC |
| | | | | K1_homology arm_ Bw | GGGTGTCGGGGCTGGCTTAGTCAAAGTATTTAGAAGCAGCT |
| pEdit_K1_kvarM | pEdit_K1 (Kan$^R$) | E/K_kvarM synthetic gene string | Editing template for site-directed *kvarM* insertion upstream of phage K1 prohead assembly protein gene (*gp168* ) | pEdit_K1_Fw* | TTATAACGCTTTTAACTTCTCTG |
| | | | | pEdit_K1_Bw* | ATAATTATGTAACTTAACACAGGA |
| | | | | K_payload_Fw | CGCAGAGAAGTTAAAAGCGTTATAAAGTACGAGGAGGTAAATATAT |
| | | | | K_kvarM_Bw | GTCCTGTGTTAAGTTACATAATTATTTATTTTTTACCAGAACCAGA |
| pEdit_K1_colE6 | pEdit_K1 (Kan$^R$) | E/K_colE6 synthetic gene string | Editing template for site-directed *colE6* insertion upstream of phage K1 prohead assembly protein gene (*gp168* ) | K_payload_Fw | CGCAGAGAAGTTAAAAGCGTTATAAAGTACGAGGAGGTAAATATAT |
| | | | | K_colE6_Bw | GTCCTGTGTTAAGTTACATAATTATTTACAGGTATTTTTTGATGTTAC |
| pEdit_K1_colE7 | pEdit_K1 (Kan$^R$) | E/K_colE7 synthetic gene string | Editing template for site-directed *colE7* insertion upstream of phage K1 prohead assembly protein gene (*gp168* ) | K_payload_Fw | CGCAGAGAAGTTAAAAGCGTTATAAAGTACGAGGAGGTAAATATAT |
| | | | | K_colE7_Bw | GTCCTGTGTTAAGTTACATAATTATTCATTTACCACGGTGGAT |
| pEdit_K1_colM | pEdit_K1 (Kan$^R$) | E/K_colM synthetic gene string | Editing template for site-directed *colM* insertion upstream of phage K1 prohead assembly protein gene (*gp168* ) | K_payload_Fw | CGCAGAGAAGTTAAAAGCGTTATAAAGTACGAGGAGGTAAATATAT |
| | | | | K_colM_Bw | GTCCTGTGTTAAGTTACATAATTATTTAACGTTTACCAGATTCTT |
| pEdit_EfS3_colE7 | $pEdit_{EfS3}$ (Em$^R$) | Efs3_colE7 synthetic gene string | Editing template for site-directed *colE7* insertion downstream of phage EfS3 *cps* | pEdit_Efs3_Fw* | CAACTAAGAAAAATTGAATAGAAAACGGA |
| | | | | pEdit_Efs3_Bw* | TATTTACCTCCTCTTAGCGTTCG |
| | | | | Efs3_payload_Fw | TCTGGAAACGTAATTAACGAACG |
| | | | | Efs3_payload_Bw | CCGTTCCCTATTCCGTTTTCTATTC |
| pEdit_EfS3_colM | $pEdit_{EfS3}$ (Em$^R$) | Efs3_colM synthetic gene string | Editing template for site-directed *colM* insertion downstream of phage EfS3 *cps* | Efs3_payload_Fw | TCTGGAAACGTAATTAACGAACG |
| | | | | Efs3_payload_Bw | CCGTTCCCTATTCCGTTTTCTATTC |
| pEdit_EfS3_kvarM | $pEdit_{EfS3}$ (Em$^R$) | Efs3_kvarM synthetic gene string | Editing template for site-directed *kvarM* insertion downstream of phage EfS3 *cps* | Efs3_payload_Fw | TCTGGAAACGTAATTAACGAACG |
| | | | | Efs3_payload_Bw | CCGTTCCCTATTCCGTTTTCTATTC |
| pEdit_EfS7_kvarM | $pEdit_{EfS7}$ (Em$^R$) | Efs3_kvarM synthetic gene string | Editing template for site-directed *kvarM* insertion downstream of phage EfS7 *cps* | pEdit_Efs7_Fw* | TAACAGCCCCCAAGAATCTTC |
| | | | | pEdit_Efs7_Bw* | CATATATATTTACCTCCTCCTACTTTCC |
| | | | | Efs7_payload_Fw | TAGGAGGAGGTAAATATATATG |
| | | | | Efs7_kvarM_Bw | GAAGATTCTTGGGGGCTGTTATTTTTTACCTGAACCTG |
| pEdit_EfS7_colE7† | $pEdit_{EfS7}$ (Em$^R$) | Efs3_colE7 synthetic gene string | Editing template for site-directed colM insertion downstream of phage EfS7 cps | Efs7_payload_Fw | TAGGAGGAGGTAAATATATATG |
| | | | | Efs7_colE7 Bw | GAAGATTCTTGGGGGCTGTTATTTACCACGGTGGATATCG |
| pEdit_EfS7_colM | $pEdit_{EfS7}$ (Em$^R$) | Efs3_colM synthetic gene string | Editing template for site-directed *colM* insertion downstream of phage EfS7 *cps* | Efs7_payload_Fw | TAGGAGGAGGTAAATATATATG |
| | | | | Efs7_colM Bw | GAAGATTCTTGGGGGCTGTTAACGTTTACCTGATTCTTTG |

**Table S4. Plasmids and primers used for phage engineering.** *, common primers for insertion of other genomic fragments into the same plasmid backbone; †, homologous recombination-based and CRISPR-Cas9-assisted engineering using plasmid pEdit_EfS7_colE7 was unsuccessful, instead the plasmid was used as a template for effector fragment production for synthetic assembly of EfS7::colE7 (Table S6).

| Name | Sequence (5'-3') | Size [bp] |
|---|---|---|
| E2_spacers | AACACGCATTGATTTGAGTCAGCTAGGAGGTGACTGAAGTATATTTTAGATGAAGATTATTTCTTAATAACTAAAAATATGGTATAATACTCTTAATAAATGCAG<br>TAATACAGGGGCTTTTCAAGACTGAAGTCTAGCTGAGACAAATAGTGCGATTACGAAATTTTTTAGACAAAAATAGTCTACGAGGTTTTAGAGCTATGCTGTTTT<br>GAATGGTCCCAAAAC**TTAATAGTCTTTTGACCGGC**GTTTTAGAGCTATGCTGTTTTGAATGGTCCCAAAAC**GTCTGATCTATAACTCCAGT**GTTTTAGAGCTAT<br>GCTGTTTTGAATGGTCCCAAAACTTCAGCACACTGAGACTTGTTGAGTTCCATGTTTTAGAGCTATGCTGTTTTGAATGGACTCCATTCAACATTGCCGATGATAA<br>CTTGAGAAAGAGGGTTAATACCAGCAGTCGGA | 452 |
| K1_spacers | AACACGCATTGATTTGAGTCAGCTAGGAGGTGACTGAAGTATATTTTAGATGAAGATTATTTCTTAATAACTAAAAATATGGTATAATACTCTTAATAAATGCAG<br>TAATACAGGGGCTTTTCAAGACTGAAGTCTAGCTGAGACAAATAGTGCGATTACGAAATTTTTTAGACAAAAATAGTCTACGAGGTTTTAGAGCTATGCTGTTTT<br>GAATGGTCCCAAAAC**CTGCGTCTACACCAACAGTC**GTTTTAGAGCTATGCTGTTTTGAATGGTCCCAAAAC**GTGAGATTCTGCCAGAGCTA**GTTTTAGAGCTA<br>TGCTGTTTTGAATGGTCCCAAAACTTCAGCACACTGAGACTTGTTGAGTTCCATGTTTTAGAGCTATGCTGTTTTGAATGGACTCCATTCAACATTGCCGATGATA<br>ACTTGAGAAAGAGGGTTAATACCAGCAGTCGGA | 452 |
| E2_homology arm | CGTAGCGTTAGTTGCTTCGCCCACAGGAGAATCATCTACATACCATTGATAAGTAGCGTTTGCGCCCGATGGCTGCGAAGCTAAAGCGGCAGTAAATTCAATAGC<br>TGTTCCGATTACTCCAGCGTCAGGGCTATTAGGAGTTACAGCTAAGGTAGTCGTTTGTGTCTTATTTTGAACTGTGATAGTTGTCGCTTCAGCCGTTTCCGGGCCT<br>CCTTCAGAAAGTGTATTTGTTGCAACTACT**TTAATAGTTTTCTGGCCTGC**AGGTCCTTTTAGTACATAACTAAAAGTTGCTTCAGCTCCATCTTGTGGAGCATCA<br>TCTACAGTCCAAGCATAGGTAATAGTTCCGCCTCCAGTTTCACCACTGGGTGTAGCAGTAAACTGCTGA**GTCTGATCGATCACCCCTGT**AGGTGTTTTAGGAGT<br>TATATCAACTGTAAAAGCCATAAGTTATCCTTATTTTAATGTTACGAAAGAAGAGTTGCGTGTTTCACGAATTAAAACTGATCCGTCACGGTTAATGTAATAAATT<br>AAGCTAAATAAAGTTTGGTGTGCTGACGCATGTTCAAAACTAGTTGGGTGAGATTTCCAATCCGGAGTTTCAGCGATCCATTGATAAATCCACCAAGGAACAGTA<br>CAGAATCCTGGATTTTTTCCAATCAACTGAAGATTCGGGCTAAAGTTTTCCGGAAGAGTAAATACAGACGGCTTTTCAGATTCAATAATCTCAGCTACAGCCTGT<br>TCGAATTTTTCTTCAACAAAAGGAGTATCTTCAATCAAAACATCGGTATTTTCAGGAATTTTATCCGTTTCTACTACTTCAATTTTAATGTCAGATTTAACTGGAG<br>AATCAATCAAAAGTGCTGCTTCTGGATTGACTTCTTCATCGTCATATT | 890 |
| E/K_kvarM | AGTACGAGGAGGTAAATATAT*ATGTCTGACACCATGATCGTTGTTGCTACCCCGACCCCGGGTTTCTCTTACGCTTCTGGTCTGACCTACGGTGGTGGTGCTTTCGCTGG*<br>*TGCTCCGGCTAACGGTCCGTCTGAAGGTCAGATCTTCTTCCAGACCGTTCTGCCGGCTTACCAGTCTCCGAACCTGTGCATCGGTCAGCTGGCTTGGATGACCGACTACAT*<br>*CAACAAAAACGGTGTTGGTAACCCGAAAACCTGGGAAGTTATCTCTCAGAACGTTCTGATCTTCTGCTCTGCTGACACCGCTCTGGTTCTGAACCCGCGTATCGCTGTTTACG*<br>*ACGGTTTCCACAAAACCAAATGGGCTCCGGCTAAATTCAACTTCAAAACCCAGTCTCAGGAAAAATTCTCTGGTAACGTTACCACCCCGATCGCTGCTTTCGGTCACTACCTG*<br>*TGGGGTGAAGGTAAACCGCGTACCGTTGACCTGTCTTCTGTTGGTCTGAAAATCCAGGCTAACCAGATCGACCCGGTTATGATCGCTGTTAAAAACAACGCTGCTGGTACCT*<br>*ACCAGATCTCTGGTAACTTCAACCGTAACACCTTCATCGACGGTGACATCCCGGGTCTGTACCTGGGTAACATCACCATGAAAACCGAAGGTACCCTGAAAATCGACGCTAA*<br>*AGGTAACTGGAACTACAACGGTGTTGTTCGTGCTTTCAACGACACCTACGACGCTAACCCGTCTACCCACCGTTCTAAATCTGCTGAAGACCTGACCACCCTGCTGCGTCTG*<br>*ACCCAGGGTACCCCGTACGAAATCCGTATCCCGGGTGAACTGAAAGTTTCTGGTTCTGGTAAAAAATAA* | 852 |
| E/K_colE7 | *AGTACGAGGAGGTAAATATAT ATGTCTGGTGGTGACGGTCGTGGTCACAACTCTGGTGCTCACAACACCGGTGGTAACATCAACGGTGGTCCGACCGGTCTGGGTGGTAAC*<br>*GGTGGTGCTTCTGACGGTTCTGGTTGGTCTTCTGAAAACAACCCGTGGGGTGGTGGTTCTGGTTCTGGTGTTCACTGGGGTGGTGGTTCAGGTCACGGTAACGGTGGTGGT*<br>*AACTCTAACTCTGGTGGTGGTTCTAACTCTTCTGTTGCTGCTCCGATGGCTTTCGGTTTCCCGGCTCTGGCTGCTCCGGGTGCTGGTACCCTGGGTATCTCTGTTTCTGGTG*<br>*AAGCTCTGTCTGCTGCTATCGCTGACATCTTCGCTGCTCTGAAAGGTCCGTTCAAATTCTCTGCTTGGGGTATCGCTCTGTACGGTATCCTGCCGTCTGAAATCGCTAAAGAC*<br>*GACCCGAACATGATGTCTAAAATCGTTACCTCTCTGCCGGCTGAAACCGTTACCAACGTTCAGGTTTCTACCCTGCCGCTGGACCAGGCTACCGTTTCTGTTACCAAACGTGT*<br>*TACCGACGTTGTTAAAGACACCCGTCAGCACATCGCTGTTGTTGCTGGTGTTCCGATGTCTGTTCCGGTTGTTAACGCTAAACCGACCCGTACCCCGGGTGTTTTCCACGCT*<br>*TCTTTCCCGGGTGTTCCGTCTCTGACCGTTTCTACCGTTAAAGGTCTGCCGGTTTCTACCACCCTGCCGCGTGGTATCACCGAAGACAAAGGTCGTACCGCTGTTCCGGCTG*<br>*GTTTCACCTTCGGTGGTGGTTCTCACGAAGCTGTTATCCGTTTCCCGAAAGAATCTGGTCAGAAACCGGTTTACGTTTCTGTTACCGACGTTCTGACCCCGGCTCAGGTTAAA*<br>*CAGCGTCAGGACGAAGAAAAACGTCTGCAGCAGGAATGGAACGACGCTCACCCGGTTGAAGTTGCTGAACGTAACTACGAACAGGCTCGTGCTGAACTGAACCAGGCTAA*<br>*CAAAGACGTTGCTCGTAACCAGGAACGTCAGGCTAAAGCTGTTCAGGTTTACAACTCTCGTAAATCTGAACTGGACGCTGCTAACAAAACCCTGGCTGACGCTAAAGCTGAA*<br>*ATCAAACAGTTCGAACGTTTCGCTCGTGAACCGATGGCTGCTGGTCACCGTATGTGGCAGATGGCTGGTCTGAAAGCTCAGCGTGCTCAGACCGACGTTAACAACAAAAAA*<br>*GCTGCTTTCGACGCTGCTGCTAAAGAAAAATCTGACGCTGACGTTGCTCTGTCTTCTGCTCTGGAACGTCGTAAACAGAAAGAAAACAAAGAAAAAGACGCTAAAGCTAAACT*<br>*GGACAAAGAATCTAAACGTAACAAACCGGGTAAAGCTACCGGTAAAGGTAAACCGGTTAACAACAAATGGCTGAACAACGCTGGTAAAGACCTGGGTTCTCCGGTTCCGGAC*<br>*CGTATCGCTAACAAACTGCGTGACAAAGAATTCAAATCTTTCGACGACTTCCGTAAAAAATTCTGGGAAGAAGTTTCTAAAGACCCGGAACTGTCTAAACAGTTCTCTCGTAAC*<br>*AACAACGACCGTATGAAAGTTGGTAAAGCTCCGAAAACCCGTACCCAGGACGTTTCTGGTAAACGTACCTCTTTCGAACTGCACCACGAAAAACCGATCTCTCAGAACGGTG*<br>*GTGTTTACGACATGGACAACATCTCTGTTGTTACCCCGAAACGTCACATCGACATCCACCGTGGTAAATGA* | 1752 |
| E/K_ec300 | AGTACGAGGAGGTAAATATAT*ATGAAAGAATACACCGCTGGTGCTGGTACCCAGCTGGCTGTTTTCCCGCTGGACGTTATCAACGTTACCCAGGGTGAAAACGGTGGTTA*<br>*CTCTCACATGGGTGCTCTGGCTATCGACTTCTCTGACGGTACCCCGCACAAACCGTACTACGCTCCGTTTGACTGCGAATGCGTATACACCGACAGCTACTCTGGTGTTGCT*<br>*TGGCAGTCTCAGAAACCGGTTAAATGCGTTGACGGTTCTGTTACCTACGTTACCCTGCTGTGCGTTCACGACAACAACTGGGCTTCTAACAAAGTTGGTGACAAAAAAGCTAA*<br>*AGGTGAAGTTATCGGTCACTCTGGTACCGCTGGTCAGGCTTCTGGTGACCACGCTCACTTCGAAGTTTCTAAAGGTAAATGGCAGGGTTGGTCTACCTCTTCTGCTGGTGTT*<br>*TACTTCATCAAAAACCCGTCTCACCTGTACGACGTTTTCTCTATCAAAAACAACGTTACCGGTAAAACCACCAAAATCATGAACGGTGGTGGTTACAACTGGCGTTCTATCGAC*<br>*TGGGACGACAAATCTGGTTCTGGTTCTGGTCCGGAACCGTCTAAACCGACCCCGTCTAAACCGTCTAACAACAAAAACAAAGAAGGTGTTGCTACCATGTACTGCCTGTACG*<br>*AACGTCCGATCAACTCTAAAACCGGTGTTCTGGAATGGAACGGTGACGCTTGGACCGTTATGTTCTGCAACGGTGTTAACTGCCGTCGTGTTTCTCACCCGGACGAAATGAA*<br>*AGTTATCGAAGACATCTACCGTAAAAACAACGGTAAAGACATCCCGTTCTACTCTCAGAAAGAATGGAACAAAAACGCTCCGTGGTACAACCGTCTGGAAACCGTTTGCCCG*<br>*GTTGTTGGTATCACCAAAAAATCTTAA* | 924 |
| immE7 | ATGGAACTGGAAAACTCTATCTCTGACTACACCGAAGCTGAATTCGTTCAGCTGCTGAAAGAAATCGAAAAAGAAAACGAAGCTGCTACCGACGACGTTCTGGA<br>CGTTCTGCTGGAACACTTCGTTAAAATCACCGAACACCCGGACGGTACCGACCTGATCTACTACCCGTCTGACAACCGTGACGACTCTCCGGAAGGTATCGTTAA<br>AGAAATCAAAGAATGGCGTGCTGCTAACGGTAAACCGGGTTTCAAACAGGGTTAA | 264 |
| immE6 | ATGGGTCTGAAACTGCACATCAACTGGTTCGACAAAACCACCGAAGAATTCAAAGGTGGTGAATACTCTAAAGACTTCGGTGACGACGGTTCTGTTATCGAACGT<br>CTGGGTATGCCGCTGAAAGACAACATCAACAACGGTTGGTTCGACGTTATCGCTGAATGGGTTCCGCTGCTGCAGCCGTACTTCAACCACCAGATCGACATCTCT<br>GACAACGAATACTTCGTTTCTTTCGACTACCGTGACGGTGACTGGTAA | 258 |
| K1_homology arm | GAGAATTAAACCATCCACCTCGTGCAAATGTTGACCCGATGCAAGCCGCAATCATTATCGAAGATATGTGGTGGAAAGGCAACGATGTTTATGGGCGTGCAAGA<br>ATCATCGAAGGAGACCACGGTCCAGGTGATAAATTAGCCGCTAACATCCGTGCTGGTTGGATTCCTGGTGTATCAAGCCGTGGGCTCGGTTCTTTAAAAGACTCA<br>GGAAAAGGATATAACATCGTACAAGAAGGCTTTCGTCT**GACTGTTGGTGTAGACGCAG**TTTGGGGTCCTAGTGCTCCGGATGCATGGGTACAACCGAAACAAA<br>TTTCGGAAAATACCTCGGCACAAGTTGCCAATAGTGCTGATGATGCTTTTATGGCTCTCGCAGAGAAGTTAAAAGCGTTATAAATAATTATGTAACTTAACACAG<br>GACATCAAAAATGCTTAAAGAACTTTTAATGACCGAAGCTAAAACTATCGATGCTTCCGTCGCTCTTGACAGTATTTTCGAATCAGTTCAGCTTTCTCCGGAAGCT<br>AAAGCTAACTTCAGCACTGTATTCGAAGCAACAGTCAAAAAACACGCAG**TAGCTCTGGCAGAATCTCAC**ATTGAAAAGATTGCTGAGAAAGCAGAAGAAAAAG<br>TAGAAGAAGAAAAAGAAAAAGCTAAAGAAGAAGCTCGCGAAGAGCTGAAAGAAGCTGCTTCTAAATACTTTGAC | 700 |
| E/K_colE6 | AGTACGAGGAGGTAAATATAT*ATGTCTGGTGGTGACGGTCGTGGTCACAACACCGGTGCTCACTCTACCTCTGGTAACATCAACGGTGGTCCGACCGGTCTGGGTGTTG*<br>*GTGGTGGTGCTTCTGACGGTTCTGGTTGGTCTTCTGAAAACAACCCGTGGGGTGGTGGTTCTGGTTCTGGTATCCACTGGGGTGGTGGTTCAGGTCACGGTAACGGTGGTG*<br>*GTAACGGTAACTCTGGTGGTGGTTCTGGCACCGGTGGTAACCTGTCTGCTGTTGCTGCTCCGGTTGCTTTCGGTTTCCCGGCTCTGTCTACCCCGGGTGCTGGTGGTCTGG*<br>*CTGTTTCTATCTCTGCTGGTGCTCTGTCTGCTGCTATCGCTGACATCATGGCTGCTCTGAAAGGTCCGTTCAAATTCGGTCTGTGGGGTGTTGCTCTGTACGGTGTTCTGCCG*<br>*TCTCAGATCGCTAAAGACGACCCGAACATGATGTCTAAAATCGTTACCTCTCTGCCGGCTGACGACATCACCGAATCTCCGGTTTCTTCTCTGCCGCTGGACAAAGCTACCGT*<br>*TAACGTTAACGTTCGTGTTGTTGACGACGTTAAAGACGAACGTCAGAACATCTCTGTTGTTTCTGGTGTTCCGATGTCTGTTCCGGTTGTTGACGCTAAACCGACCGAACGTC*<br>*CGGGTGTTTTCACCGCTTCTATCCCGGGTGCTCCGGTTCTGAACATCTCTGTTAACAACTCTACCCCGGCTGTTCAGACCCTGTCTCCGGGTGTTACCAACAACACCGACAA*<br>*AGACGTTCGTCCGGCTGGTTTCACCCAGGGTGGTAACACCCGTGACGCTGTTATCCGTTTCCCGAAAGACTCTGGTCACAACGCTGTTTACGTTTCTGTTTCTGACGTTCTGT*<br>*CTCCGGACCAGGTTAAACAGCGTCAGGACGAAGAAAACCGTCGTCAGCAGGAATGGGACGCTACCCACCCGGTTGAAGCTGCTGAACGTAACTACGAACGTGCTCGTGCT*<br>*GAACTGAACCAGGCTAACGAAGACGTTGCTCGTAACCAGGAACGTCAGGCTAAAGCTGTTCAGGTTTACAACTCTCGTAAATCTGAACTGGACGCTGCTAACAAAACCCTGG*<br>*CTGACGCTATCGCTGAAATCAAACAGTTCAACCGTTTCGCTCACGACCCGATGGCTGGTGGTCACCGTATGTGGCAGATGGCTGGTCTGAAAGCTCAGCGTGCTCAGACCG*<br>*ACGTTAACAACAAACAGGCTGCTTTCGACGCTGCTGCTAAAGAAAAATCTGACGCTGACGCTGCTCTGTCTTCTGCAATGGAATCTCGTAAAAAAAAAGAAGACAAAAAACGT*<br>*TCTGCTGAAAACAAACTGAACGAAGAAAAAAACAAACCGCGTAAAGGTGTTAAAGACTACGGTCACGACTACCACCCGGACCCGAAAACCGAAGACATCAAAGGTCTGGGTG*<br>*AACTGAAAGAAGGTAAACCGAAAACCCCGAAACAGGGTGGTGGTGGTAAACGTGCTCGTTGGTACGGTGACAAAGGTCGTAAAATCTACGAATGGGACTCTCAGCACGGTG*<br>*AACTGGAAGGTTACCGTGCTTCTGACGGTCAGCACCTGGGTTCTTTCGAACCGAAAACCGGTAACCAGCTGAAAGGTCCGGACCCGAAACGTAACATCAAAAAATACCTGTA*<br>*A* | 1677 |
| E/K_colM | AGTACGAGGAGGTAAATATAT*ATGGAAACCCTGACCGTTCACGCTCCGTCTCCGTCTACCAACCTGCCGTCTTACGGTAACGGTGCTTTCTCTCTGTCTGCTCCGCACGTT*<br>*CCGGGTGCTGGTCCGCTGCTGGTTCAGGTTGTTTACTCTTTCTTCCAGTCTCCGAACATGTGCCTGCAGGCTCTGACCCAGCTGGAAGACTACATCAAAAAACACGGTGCTT*<br>*CTAACCCGCTGACCCTGCAGATCATCTCTACCAACATCGGTTACTTCTGCAACGCTGACCGTAACCTGGTTCTGCACCCGGGTATCTCTGTTTACGACGCTTACCACTTCGCT*<br>*AAACCGGCTCCGTCTCAGTACGACTACCGTTCTATGAACATGAAACAGATGTCTGGTAACGTTACCACCCCGATCGTTGCTCTGGCTCACTACCTGTGGGGTAACGGTGCTG*<br>*AACGTTCTGTTAACATCGCTAACATCGGTCTGAAAATCTCTCCGATGAAAATCAACCAGATCAAAGACATCATCAAATCTGGTGTTGTTGGTACCTTCCCGGTTTCTACCAAATT*<br>*CACCCACGCTACCGGTGACTACAACGTTATCACCGGTGCTTACCTGGGTAACATCACCCTGAAAACCGAAGGTACCCTGACCATCTCTGCTAACGGTTCTTGGACCTACAAC*<br>*GGTGTTGTTCGTTCTTACGACGACAAATACGACTTCAACGCTTCTACCCACCGTGGTATCATCGGTGAATCTCTGACCCGTCTGGGTGCTATGTTCTCTGGTAAAGAATACCA*<br>*GATCCTGCTGCCGGGTGAAATCCACATCAAAGAATCTGGTAAACGTTAA* | 837 |
| Efs3_colE7 | CATTCCATCTGGAAACGTAATTAACGAACGCTAAGAGGAGGTAAATATAT*ATGTCAGGTGGTGATGGTCGTGGTCACAACTCAGGTGCTCACAACACAGGTGGTAACA*<br>*TCAACGGTGGTCCAACAGGTTTAGGTGGTAACGGTGGTGCTTCAGATGGTTCAGGTTGGTCATCAGAAAACAACCCATGGGGTGGTGGTTCAGGTTCAGGTGTTCACTGGG*<br>*GTGGTGGTTCCGGTCACGGTAACGGTGGTGGTAACTCAAACTCAGGTGGTGGTTCAAACTCATCAGTTGCTGCTCCAATGGCTTTCGGTTTCCCAGCTTTAGCTGCTCCAGG*<br>*TGCTGGTACATTAGGTATCTCAGTTTCAGGTGAAGCTTTATCAGCTGCTATCGCTGATATCTTCGCTGCTTTAAAAGGTCCATTCAAATTCTCAGCTTGGGGTATCGCTTTATAC*<br>*GGTATCTTACCATCAGAAATCGCTAAAGATGATCCAAACATGATGTCAAAAATCGTTACATCATTACCAGCTGAAACAGTTACAAACGTTCAAGTTTCAACATTACCATTAGATC*<br>*AAGCTACAGTTTCAGTTACAAAACGTGTTACAGATGTTGTTAAAGATACACGTCAACACATCGCTGTTGTTGCTGGTGTTCCAATGTCAGTTCCAGTTGTTAACGCTAAACCAA*<br>*CACGTACACCAGGTGTTTTCCACGCTTCATTCCCAGGTGTTCCATCATTAACAGTTTCAACAGTTAAAGGTTTACCAGTTTCAACAACATTACCACGTGGTATCACAGAAGATAA*<br>*AGGTCGTACAGCTGTTCCAGCTGGTTTCACATTCGGTGGTGGTTCACACGAAGCTGTTATCCGTTTCCCAAAAGAATCAGGTCAAAAACCAGTTTACGTTTCAGTTACAGATG*<br>*TTTTAACACCAGCTCAAGTTAAACAACGTCAAGATGAAGAAAAACGTTTACAACAAGAATGGAACGATGCTCACCCAGTTGAAGTTGCTGAACGTAACTACGAACAAGCTCGT*<br>*GCTGAATTAAACCAAGCTAACAAAGATGTTGCTCGTAACCAAGAACGTCAAGCTAAAGCTGTTCAAGTTTACAACTCACGTAAATCAGAATTAGATGCTGCTAACAAAACATTA*<br>*GCTGATGCTAAAGCTGAAATCAAACAATTCGAACGTTTCGCTCGTGAACCAATGGCTGCTGGTCACCGTATGTGGCAAATGGCTGGTTTAAAAGCTCAACGTGCTCAAACAGA*<br>*TGTTAACAACAAAAAAGCTGCTTTCGATGCTGCTGCTAAAGAAAAATCAGATGCTGATGTTGCTTTATCATCAGCTTTAGAACGTCGTAAACAAAAAGAAAACAAAGAAAAAGAT*<br>*GCTAAAGCTAAATTAGATAAAGAATCAAAACGTAACAAACCAGGTAAAGCTACAGGTAAAGGTAAACCAGTTAACAACAAATGGTTAAACAACGCTGGTAAAGATTTAGGTTCA*<br>*CCAGTTCCAGATCGTATCGCTAACAAATTACGTGATAAAGAATTCAAATCATTCGATGATTTCCGTAAAAAATTCTGGGAAGAAGTTTCAAAAGATCCAGAATTATCAAAACAATT*<br>*CTCACGTAACAACAACGATCGTATGAAAGTTGGTAAAGCTCCAAAAACACGTACACAAGATGTATCAGGTAAACGTACATCATTCGAATTACACCACGAAAAACCAATCTCACA*<br>*AAACGGTGGTGTTTACGATATGGATAACATCTCAGTTGTTACACCAAAACGTCACATCGATATCCACCGTGGTAAATAA* TCTTAGTTCAACTAAGAAAAATTGAATAGAAA<br>ACGGAATAGGGAACGGTCTAACTG | 1837 |
| Efs3_colM | CATTCCATCTGGAAACGTAATTAACGAACGCTAAGAGGAGGTAAATATAT*ATGGAAACATTAACAGTTCACGCTCCATCACCATCAACAAACTTACCATCATACGGTAAC*<br>*GGTGCTTTCTCATTATCAGCTCCACACGTTCCAGGTGCTGGTCCATTATTAGTTCAAGTTGTTTACTCATTCTTCCAATCACCAAACATGTGTTTACAAGCTTTAACACAATTAGA*<br>*AGATTACATCAAAAAACACGGTGCTTCAAACCCATTAACATTACAAATCATCTCAACAAACATCGGTTACTTCTGTAACGCTGATCGTAACTTAGTTTTACACCCAGGTATCTCA*<br>*GTTTACGATGCTTACCACTTCGCTAAACCAGCTCCATCACAATACGATTACCGTTCAATGAACATGAAACAAATGTCAGGTAACGTTACAACACCAATCGTTGCTTTAGCTCACT*<br>*ACTTATGGGGTAACGGTGCTGAACGTTCAGTTAACATCGCTAACATCGGTTTAAAAATCTCACCAATGAAAATCAACCAAATCAAAGATATCATCAAATCAGGTGTTGTTGGTA*<br>*CATTCCCAGTTTCAACAAAATTCACACACGCTACAGGTGATTACAACGTTATCACAGGTGCTTACTTAGGTAACATCACATTAAAAACAGAAGGTACATTAACAATCTCAGCTAA*<br>*CGGTTCATGGACATACAACGGTGTTGTTCGTTCATACGATGATAAATACGATTTCAACGCTTCAACACACCGTGGTATCATCGGTGAATCATTAACACGTTTAGGTGCTATGTT*<br>*CTCAGGTAAAGAATACCAAATCTTATTACCAGGTGAAATCCACATCAAAGAATCAGGTAAACGTTAA* TCTTAGTTCAACTAAGAAAAATTGAATAGAAAACGGAATAGGG<br>AACGGTCTAACTG | 922 |
| Efs3_kvarM | CATTCCATCTGGAAACGTAATTAACGAACGCTAAGAGGAGGTAAATATAT*ATGTCAGATACAATGATCGTTGTTGCTACACCAACACCAGGTTTCTCATACGCTTCAGG*<br>*TTTAACATACGGTGGTGGTGCTTTCGCTGGTGCTCCAGCTAACGGTCCATCAGAAGGTCAAATCTTCTTCCAAACAGTTTTACCAGCTTACCAATCACCAAACTTATGTATCGG*<br>*TCAATTAGCTTGGATGACAGATTACATCAACAAAAACGGTGTTGGTAACCCAAAAACATGGGAAGTTATCTCACAAAACGTTTTAATCTTCTGTTCAGCTGATACAGCTTTAGTT*<br>*TTAAACCCACGTATCGCTGTTTACGATGGTTTCCACAAAACAAAATGGGCTCCAGCTAAATTCAACTTCAAAACACAATCACAAGAAAAATTCTCAGGTAACGTTACAACACCAA*<br>*TCGCTGCTTTCGGTCACTACTTATGGGGTGAAGGTAAACCACGTACAGTTGATTTATCATCAGTTGGTTTAAAAATCCAAGCTAACCAAATCGATCCAGTTATGATCGCTGTTA*<br>*AAAACAACGCTGCTGGTACATACCAAATCTCAGGTAACTTCAACCGTAACACATTCATCGATGGTGATATCCCAGGTTTATACTTAGGTAACATCACAATGAAAACAGAAGGTA*<br>*CATTAAAAATCGATGCTAAAGGTAACTGGAACTACAACGGTGTTGTTCGTGCTTTCAACGATACATACGATGCTAACCCATCAACACACCGTTCAAAATCAGCTGAAGATTTAA*<br>*CAACATTATTACGTTTAACACAAGGTACACCATACGAAATCCGTATCCCAGGTGAATTAAAAGTTTCAGGTTCAGGTAAAAAATAA* TCTTAGTTCAACTAAGAAAAATTGAAT<br>AGAAAACGGAATAGGGAACGGTCTAACTG | 937 |

**Table S5. Synthetic DNA strings used for pSelect and pEdit plasmid generation.** Underlined = direct repeat; bold = spacer; Red = mutated PAM or seed sequence; Green = RBS; Italics = coding sequence of inserted effector gene

| HEPT synthetic phage | Fragment ID | Template | Primers | Primer Sequence (5'-3') |
|---|---|---|---|---|
| CM001::*ec300* | CM001 F1 | CM001 gDNA | CM001 F1.1 Fw | TCAACGCTTGACAGCCGCA |
| | | | CM001 F1.1 Bw | TTACTTGTCCGCGTCGGCG |
| | CM001_ec300 | E/K_ec300 synthetic gene string | CM001_ec300 Fw | GGCAATCGCCGACGCGGACAAGTAAAGTACGAGGAGGTAAATATAT |
| | | | CM001_ec300 Bw | CCCCTTTGTTTTTACTCCAACCGTATTAAGATTTTTTGGTGATACC |
| | CM001 F1.2 | CM001 gDNA | CM001 F1.2 Fw | TACGGTTGGAGTAAAAACAAAGGG |
| | | | CM001 F1.2 Bw | TCGGCCTCAGCTTCGTAATAA |
| | CM001 F2 | CM001 gDNA | CM001 F2 Fw | AAGGATTAAATAATGAAACTTTCTGATTT |
| | | | CM001 F2 Bw | TTATTTATCTTCTAGTGCTGCCA |
| | CM001 F3 | CM001 gDNA | CM001 F3 Fw | GTTCTTCATAGAGATTGCCTATC |
| | | | CM001 F3 Bw | AGTACAAATCATCAAAGTAAGCA |
| | CM001 F4 | CM001 gDNA | CM001 F4 Fw | ACTATTAACTTTGTCATTGTATAATCCTTGT |
| | | | CM001 F4 Bw | ACCCTGCAAATCGTACTGGT |
| EfS7::*colE7* | EfS7 F1.1 | EfS7 gDNA | EfS7 F1.1 Fw | AAAATCCTCTATAAGGCGTCC |
| | | | EfS7 F1.1 Bw | ACTTGAGCATCAATAACCCAC |
| | EfS7_colE7 | pEdit_EfS7_colE7 | EfS7 colE7 i Fw | AAATGACTGATAGCTACGAGTG |
| | | | EfS7 colE7 i Bw | TTTGCTTCCGTCAGAAGC |
| | EfS7 F2.1 | EfS7 gDNA | EfS7 F2.1 Fw | TTCTCAAAAGACTATGTCTCTAGC |
| | | | EfS7 F2.1 Bw | AACCCTTGCAAACTTCTTACC |
| | EfS7 F3 | EfS7 gDNA | EfS7 F3 Fw | ACTTGCCTCCTGAAACTTGG |
| | | | EfS7 F3 Bw | TCCTTTAGTGTCATTATCAGTGC |
| | EfS7 F4 | EfS7 gDNA | EfS7 F4 Fw | CGACAACATCATCATAGGCACT |
| | | | EfS7 F4 Bw | GTTCATCATAACCTACGTGACC |
| | EfS7 F4 | EfS7 gDNA | EfS7 F5 Fw | CATAAATCCATTCTAAGAGGTCACG |
| | | | EfS7 F5 Bw | CTTGTCGGGAAGTGTGTC |

**Table S6. Primers and templates used for synthetic HEPTs construction.**
